# Multi-omic profiling of intraductal papillary neoplasms of the pancreas reveals distinct expression patterns and potential markers of progression

**DOI:** 10.1101/2024.07.07.602385

**Authors:** Yuefan Wang, T. Mamie Lih, Jae W. Lee, Takao Ohtsuka, Yuto Hozaka, Mari Mino-Kenudson, N. Volkan Adsay, Claudio Luchini, Aldo Scarpa, Ajay V. Maker, Grace E. Kim, Jorge Paulino, Lijun Chen, Liyuan Jiao, Zhenyu Sun, Davina Goodman, Michael J. Pflüger, Nicholas J. Roberts, Hanno Matthaei, Laura D. Wood, Toru Furukawa, Hui Zhang, Ralph H. Hruban

**Affiliations:** Department of Pathology, Sol Goldman Pancreatic Cancer Research Center, Johns Hopkins University School of Medicine, Baltimore, MD, USA; Department of Digestive Surgery, Kagoshima University, Kagoshima, Japan; Department of Pathology, Massachusetts General Hospital & Harvard Medical School, Boston, MA, USA; Department of Pathology, Koç University School of Medicine, Istanbul, Turkey; Department of Diagnostics and Public Health, Section of Pathology, ARC-Net Research Center, University and Hospital Trust of Verona, Verona, Italy; Division of Surgical Oncology, Department of Surgery, University of California San Francisco, San Francisco, CA, USA; Department of Pathology, University of California San Francisco, San Francisco, CA, USA; Department of Surgery, Hospital da Luz, Lisbon, Portugal; Department of Oncology, Sol Goldman Pancreatic Cancer Research Center, Sidney Kimmel Cancer Center, Johns Hopkins University School of Medicine, Baltimore, MD, USA; Department of Surgery, University Hospital of Bonn, Bonn, Germany; Department of Investigative Pathology, Tohoku University Graduate School of Medicine, Sendai, Japan

## Abstract

In order to advance our understanding of precancers in the pancreas, 69 pancreatic intraductal papillary neoplasms (IPNs), including 64 intraductal papillary mucinous neoplasms (IPMNs) and 5 intraductal oncocytic papillary neoplasms (IOPNs), 32 pancreatic cyst fluid samples, 104 invasive pancreatic ductal adenocarcinomas (PDACs), 43 normal adjacent tissues (NATs), and 76 macro-dissected normal pancreatic ducts (NDs) were analyzed by mass spectrometry. A total of 10,246 proteins and 22,284 glycopeptides were identified in all tissue samples, and 756 proteins with more than 1.5-fold increase in abundance in IPMNs relative to NDs were identified, 45% of which were also identified in cyst fluids. The over-expression of selected proteins was validated by immunolabeling. Proteins and glycoproteins overexpressed in IPMNs included those involved in glycan biosynthesis and the immune system. In addition, multiomics clustering identified two subtypes of IPMNs. This study provides a foundation for understanding tumor progression and targets for earlier detection and therapies.

**Significance:** This multilevel characterization of intraductal papillary neoplasms of the pancreas provides a foundation for understanding the changes in protein and glycoprotein expression during the progression from normal duct to intraductal papillary neoplasm, and to invasive pancreatic carcinoma, providing a foundation for informed approaches to earlier detection and treatment.

## Introduction

Pancreatic cancer is an aggressive disease with a five-year survival rate of only 13% ^1^. In contrast to many other cancers, deaths from pancreatic cancer are increasing, and it has been predicted that pancreatic cancer will become the second leading cause of cancer death in the United States and in parts of Europe by the year 2030 ^2-5^. Early detection has dramatically reduced deaths from other cancer types including colorectal cancer, and the survival rate of patients with early-stage pancreatic cancer is significantly higher than that of those with advanced-stage disease ^6,7^. This supports the premise that earlier detection of invasive cancer is a crucial approach to reduce deaths from pancreatic cancer ^6,7^. However, even small invasive pancreatic cancers have proven deadly, implying that screening for non-invasive precursor lesions, such as intraductal papillary neoplasms (IPNs), is an effective approach to prevent deaths from pancreatic cancer ^8,9^.

IPNs, including intraductal papillary mucinous neoplasms (IPMNs), intraductal oncocytic papillary neoplasms (IOPNs) and intraductal tubulopapillary neoplasms (ITPNs), are macroscopic non-invasive precursor lesions, some of which progress to invasive pancreatic cancer ^10-14^. IPNs form cysts in the pancreas, and these cysts are detectable with currently available imaging technologies ^15^. Pancreatic cysts have a variety of causes and are often misclassified clinically, sometimes leading to incorrect preoperative diagnoses and unnecessary surgery ^16,17^. Even when correctly diagnosed, the opportunities to cure a precancer through surgical resection need to be balanced with the risk of significant surgical complications ^18,19^. New approaches are needed to accurately diagnose and stratify the risk of progression of IPNs^20,21.^

Proteins and glycoproteins have proven to be useful clinical biomarkers for classifying cyst-forming neoplasms in the pancreas. For example, vascular endothelial growth factor (VEGF) is often overexpressed in benign serous cystadenomas, while elevated levels of carcinoembryonic antigen (CEA) and cancer antigen 19-9 (CA-19.9) in cyst fluids suggest mucin producing cystic neoplasms, such as IPMNs or mucinous cystic neoplasms, and at very high levels, they can indicate a risk of malignancy ^22-26^. However, the available biomarkers for classifying cyst types are imperfect, and a better understanding of pancreatic cancer biology could provide more sensitive and specific markers ^27-29^.

Modern mass spectrometry has proven a powerful tool in the characterization of the patterns of protein and glycoprotein expression in a variety of tumor types, and it has led to the discovery of new biomarkers of tumor subtypes, novel cellular pathways, and new markers of prognosis ^30-38^. For example, an integrated proteogenomic analysis of 140 pancreatic ductal adenocarcinomas (PDACs) revealed associations between somatic mutations and changes in protein expression and helped delineate unique molecularly defined subtypes of PDAC ^33^.

In this study we used proteomics and glycoproteomics to characterize a series of histologically and genetically well-characterized surgically resected IPNs, including IPMNs and IOPNs. As described previously, the use of macro-dissected normal pancreatic ducts allowed us to compare the neoplastic cells of the IPNs to the appropriate non-neoplastic cells (neoplastic cells with ductal differentiation to non-neoplastic ductal cells) ^33^. The inclusion of paired cyst fluid samples aspirated from the neoplasms allowed us to determine which proteins and glycoproteins expressed in the neoplastic cells of the IPNs are detectable in cyst fluids, a clinically obtainable sample type ^39^. Finally, we reanalyzed a series of previously reported well-characterized PDAC samples ^33^ using the same platform that the IPN samples were analyzed on, allowing us to compare protein expression patterns in precursor lesions to those in invasive cancers of the same organ.

## Results

### Landscape of the intraductal papillary neoplasm cohort

Our comprehensive multi-level analysis included 69 IPNs (64 IPMNs and 5 IOPNs), 76 macro-dissected grossly normal duct tissues (NDs, with 32 of the 76 NDs coming from the same resection specimens as the IPNs), and 32 cyst fluid samples (featuring 22 from matched IPMN patients, 3 matched IOPN patients, 2 from unrelated IPMN patients, and 5 other cyst samples) (Fig. 1A). The use of macro-dissected normal pancreatic ducts as controls was critical, as IPNs have ductal differentiation, while bulk normal pancreas is composed primarily of acinar cells.

**Figure 1.**
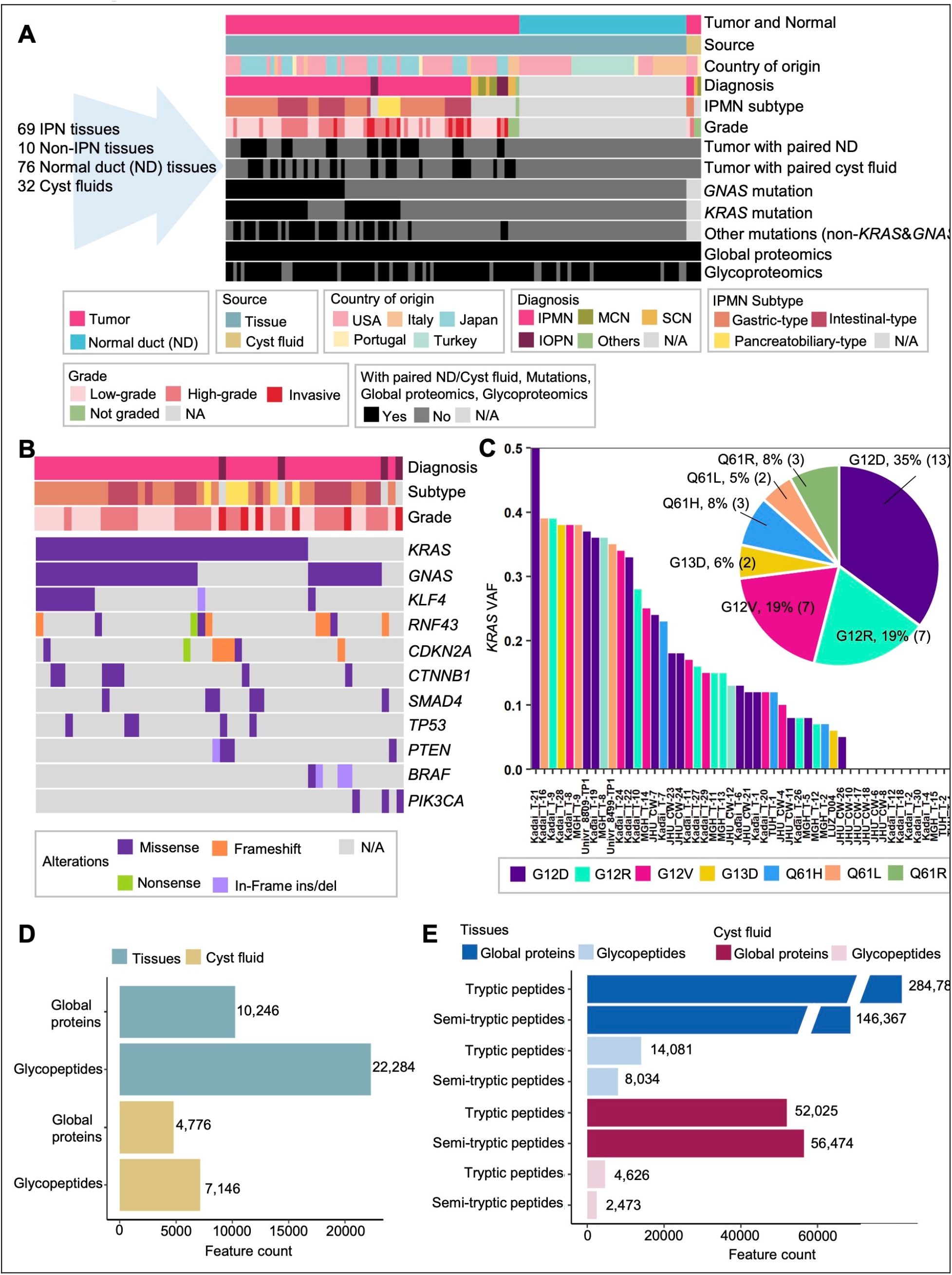
Landscape of the cohort. **A**. Distribution of the cohort according to sample types, country of origin, histologic subtypes, histologic grade, and available data types. **B**. Mutation landscape of 50 IPNs (46 IPMNs and 4 IOPNs). **C**. Distribution of *KRAS* VAF and *KRAS* hotspot mutations in 37 IPNs (35 IPMNs and 2 IOPNs). **D**. Total global proteins and glycopeptides identified from the global proteomics and glycoproteomics of tissues and cystic fluids of the entire cohort. **E**. Identification of fully-tryptic and semi-tryptic peptides from the global proteomics and glycoproteomics of tissues and cystic fluids of the entire cohort.

Fresh lesional tissues and normal ductal tissue samples were collected from patients who underwent pancreas surgery in five different countries (Fig. 1A). Corresponding clinical data and centralized pathology review by one of us (T.F.) are summarized in Supplementary Table S1A. The age and sex of these patients from whom the IPNs were collected are similar to those reported in the general population of patients who have undergone surgical resection for IPNs^40,41^. Similarly, the genes identified as somatically mutated in these neoplasms (Fig. 1B, Supplementary Table S1B-S1D) are similar to those previously reported for IPNs of the pancreas^28,42,43^. In particular, 37 of 69 IPNs (54%) harbored a *KRAS* mutation, and 32 of the 69 (47%) aGNAS mutation. Moreover, among 33 IPNs with a *KRAS* mutation variant allele fraction (VAF) > 0.075, the majority harbored one of the common hot-spot *KRAS* mutations of G12D (35%), G12R (19%), or G12V (19%) (Fig. 1C). Similarly, among the IPNs with a GNAS mutation with a VAF >0.075, the majority harbored one of the *GNAS* hotspot mutations of R201C (50%) or R201H (41%) (Supplementary Figs. S1A and S1B). Specifically, GNAS R201C was mostly observed in intestinal-type IPMNs, while gastric-type of IPMNs more commonly harbored *GNAS* R201H mutation, with none of the pancreatobiliary-type IPMNs having hotspot *GNAS* mutations (Supplementary Fig. S1C).

### Application of mass spectrometry to intraductal papillary neoplasms

Utilizing trapped ion mobility spectrometry (TIMS) high-throughput 4D-proteomics to conduct data-independent acquisition mass spectrometry (DIA-MS) for both proteomics and glycoproteomics analysis ^44-46^, we identified and quantified a total of 10,246 proteins and 22,284 glycopeptides in all tissue samples, as well as 4,776 proteins and 7,146 glycopeptides in the pancreatic cyst fluid samples (Fig. 1D). The median number of identified proteins in tissues was 8,041, and the median number of glycopeptides was 8,100. For cyst fluids, the median identification numbers for proteins and glycopeptides were 1,826 and 1,591, respectively (Supplementary Fig. S1D). Intriguingly, a multitude of semi-tryptic (non-tryptic end) peptides were detected (Fig. 1E), indicating the impact of pancreatic digestive enzymes on protein and glycoprotein integrity.

### The protein landscape of intraductal papillary mucinous neoplasms

Given their unique direction of differentiation^47^, the 5 IOPNs were not included in the following analyses. Comparing the proteins identified in the 64 IPMNs to those in 76 NDs revealed that 756 proteins were significantly up-regulated by more than 1.5-fold and 438 proteins were down-regulated by more than 1.5-fold in the IPMNs (Fig. 2A, Supplementary Table S2A). As implied by the inclusion of “mucinous” in the name of IPMNs, glycans and polysaccharides were significantly upregulated in these neoplasms. The IPMNs that harbored *KRAS* and/or *GNAS* hotspot mutations exhibited particularly elevated glycosylation and glycoprotein processing proteins (Supplementary Figs. S2A and Table S2B-S2D). Kyoto Encyclopedia of Genes and Genomes (KEGG) pathway enrichment analysis showed a significant enrichment of the IPMN proteome in glycan biosynthesis and metabolism pathways,particularly mucin type O-glycan biosynthesis (Fig. 2B and Supplementary Table S2E) ^48-50^. Additionally, immune pathways related to host defense were identified, where the immune system recognizes and eliminates antibody-tagged pathogens ^51^. Similar patterns of protein expression were observed when only the 46 IPMNs with a somatic mutation were considered (Supplementary Fig. S2B and Table S2F). The proteins expressed in the IOPNs are separately listed in Supplementary Table S2G.

**Figure 2.**
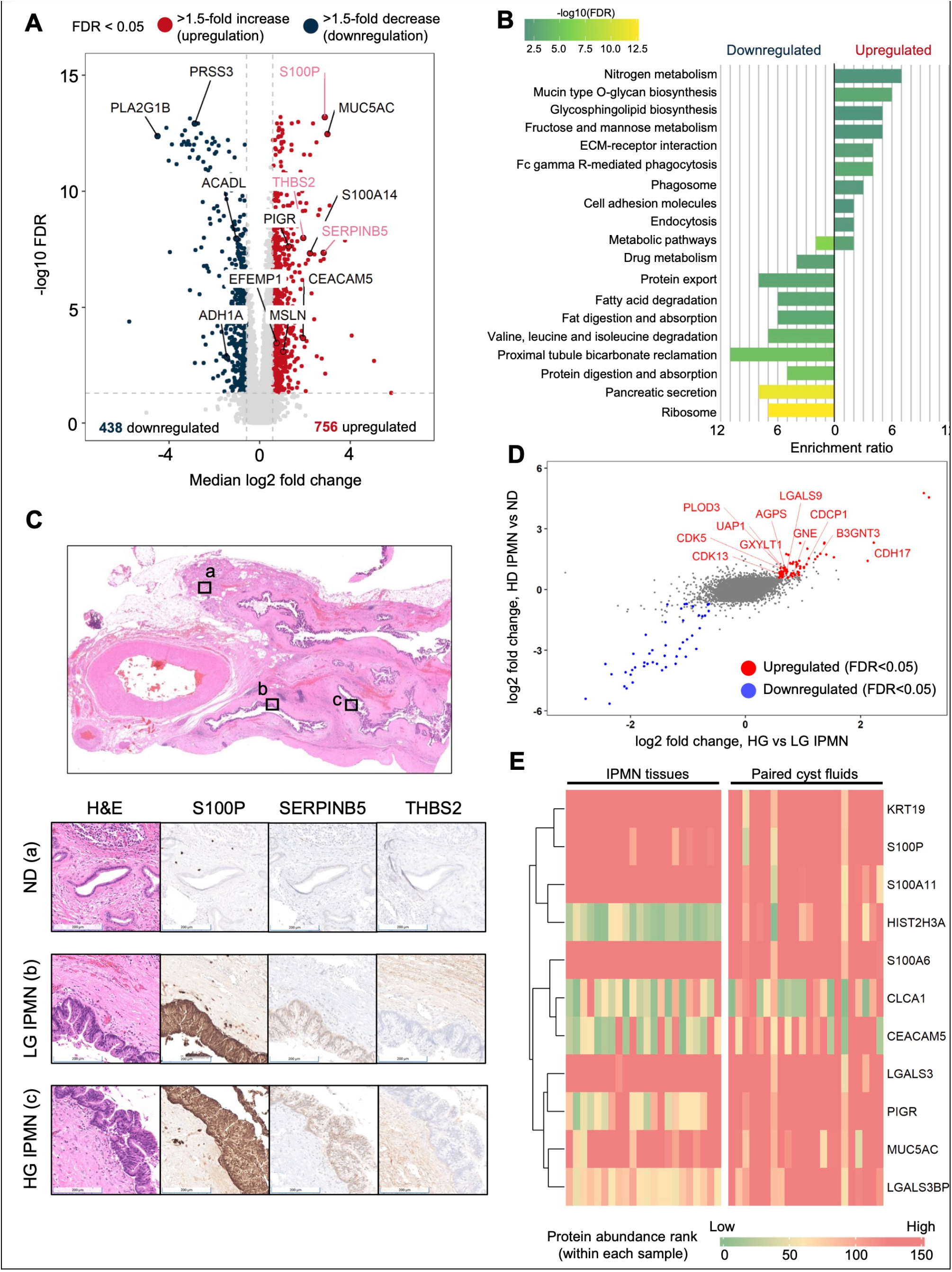
Identification of IPMN- and grade-associated proteins and detection potential using cyst fluid **A**. Comparison of 64 IPMNs and 76 normal duct tissues (NDs). Proteins with immunochemistry labeling are in pink. **B**. KEGG pathway enriched by using upregulated and downregulated proteins in IPMN tissues relative to NDs. **C**. IHC labeling results of S100P, SERPINB5, and THBS2 in ND, low-grade (LG) IPMN, and high-grade (HG) IPMN. **D**. Significantly upregulated (red) and downregulated (blue) proteins in HG IPMNs relative to LG IPMNs and NDs. Proteins associate with glycosylation/cancer progression are named in red. **E**. Expression profile comparison between IPMNs and paired cyst fluids (n=22) of some differentially expressed tissue proteins that were found as overexpressed in IPMNs or HG IPMNs in comparison of either NDs or LG IPMNs.

Notably, 88 of the 756 proteins overexpressed in the IPMNs were also reported as upregulated in PDACs in our previously published study ^33^ (Supplementary Table S2H). Some proteins previously reported as overexpressed in low-stage PDAC, such as S100P, SERPINB5, and THBS2, were also upregulated in the IPMNs relative to NDs (highlighted in pink in Fig. 2A) ^33^. Conversely, proteins highly abundant in NDs were associated with normal pancreatic functions, including protein and fatty acid digestion and absorption, as well as pancreatic secretion (Fig.2B).

### Localization of protein expression

Tissue samples of intraductal neoplasms and invasive cancers contain a mixture of neoplastic and non-neoplastic cells ^52,53^. Proteins overexpressed by the neoplastic cells are more likely to be released into cyst fluid than those expressed in the stroma. Therefore, we utilized a previously published database that identified genes differentially expressed in the neoplastic cells and stroma of PDAC to define the most likely compartment for each overexpressed protein identified in the IPMN tissues^54^. As shown in Supplementary Table S2I, transcripts such as S100P and SERPINB5 were previously shown to be elevated in PDAC neoplastic cells, whereas transcripts such as thrombospondin 2 (THBS2), and polymeric immunoglobulin receptor (PIGR) were previously identified as elevated in the stromal compartment. The thrombospondins function in cell-matrix interactions, and PIGR has been reported to be a marker of immunologically “hot” tumor nests ^54,55^.

We next validated the expression of selected proteins by immunolabeling. This not only allowed us to validate the overexpression of these proteins but also allowed us to determine the compartments in which they were expressed. As shown in Fig. 2C and Supplementary Table S2J, as expected, antibodies to S100P and SERPINB5 demonstrated that these proteins were overexpressed by the neoplastic cells, while antibodies to THBS2 labeled the stromal cells.

### Protein expression by tumor grade/histology subtype

It is clinically important to distinguish IPMNs with low-grade dysplasia from those with high-grade dysplasia, and from IPMNs with an associated invasive carcinoma^56^. Those with low-grade dysplasia can be safely followed, while the presence of high-grade dysplasia or IPMNs with an associated invasive carcinoma (the term “high-grade IPMN” in following analyses refers to both high-grade dysplasia and IPMN with an associated invasive carcinoma together) typically warrants surgical resection^56,57^. There were 68 proteins overexpressed in high-grade IPMNs relative to those with low-grade dysplasia (Supplementary Table S2K). These included proteins related to glycosylation biosynthesis, such as PLOD3, GXYLT1, UAP1, GNE, and B3GNT3. Among these glycan synthetic proteins, GNE (Glucosamine (UDP-N-acetyl)-2-epimerase/N-acetylmannosamine kinase) is a key enzyme required for the modification of sialic acid on glycoproteins, a crucial modification for the stability and recognition of glycoproteins, and implicated in tumor progression^58^. Other upregulated proteins such as CDH17, AGPS, CDK5, LGALS9, CDCP1, and CDK13 are involved in pathways and networks that influence tumor invasion and progression. To identify proteins associated with high-grade IPMNs with progression risk, we found proteins overexpressed in both high-grade IPMNs relative to low-grade IPMNs and high-grade IPMNs relative to NDs. (Supplementary Table S2K, highlighted in red in Fig. 2D). Some of these upregulated proteins, such as galectin 9 (LGALS9), multifunctional procollagen lysine hydroxylase and glycosyltransferase LH3 (PLOD3), and insulin receptor substrate (IRS2), have been previously reported as upregulated in PDAC and have been reported to play a role in cancer progression ^59-64^.

Additionally, different histological subtypes of IPMNs show distinct protein expression patterns. In gastric-type IPMNs, 47 proteins were upregulated compared to other IPMNs, with 23 of these proteins also showing overexpression in low-grade IPMNs compared to high-grade IPMNs (Supplementary Table S2L). For the intestinal-type IPMNs, 57 proteins were upregulated, 14 of which were also upregulated in high-grade IPMNs compared to low-grade IPMNs (Supplementary Table S2M). The expression of the digestive enzyme carboxypeptidase A (CPA) gradually decreased from gastric-type to intestinal-type to pancreatobiliary-type (Supplementary Fig. S2C). In contrast, proteins such as CEACAM5, MUC2, GNE, and CDH17 were overexpressed in the intestinal-type compared to other IPMNs (Supplementary Fig. S2D). MUC2 involves in the key process of intestinal-type IPMN differentiation and progression^65^. These results suggest potential biomarkers for tumor grade/histology subtype and progression.

### Proteins identified in the IPMNs can be detected in pancreatic cyst fluid

Cyst fluid can be sampled endoscopically, providing a simple way to classify cysts clinically if the proteins expressed by the neoplastic cells are released into the cyst contents^22-28^. Therefore, we compared, among the proteins upregulated in IPMNs, the patterns of protein expression in IPMNs with those in paired cyst fluid samples (n=22) (Supplementary Fig. 2E). As shown in Fig. 2E, a number of proteins upregulated in IPMNs were also upregulated in cyst fluid samples. Members of the S100 protein family, including S100A6, S100A11, and S100P, were highly expressed in IPMNs and abundant in the cyst fluid samples, as was one of the major carrier proteins for CA-19-9, MUC5AC. To allow for a more complete exploration of the clinical utility of a number of markers in cyst fluid, we list commonly identified and quantified proteins in tissue and cyst fluid samples (Supplementary Table S2N).

### The glycoprotein landscape of intraductal papillary mucinous neoplasms

Comparison of the glycopeptides identified in the 53 IPMNs (among 64 IPMNs, 11 lacked enough digested global peptides for glycopeptide enrichment and the 5 IOPNs were excluded in these analyses) with available glycopeptide data to those identified in the 66 NDs (among 76 NDs, 10 lacked enough digested global peptides for glycopeptide enrichment) revealed that 2,327 glycopeptides were significantly upregulated more than 1.5-fold and 1,270 glycopeptides were downregulated more than 1.5-fold in the IPMNs (Fig. 3A, Supplementary Table S3A).

**Figure 3.**
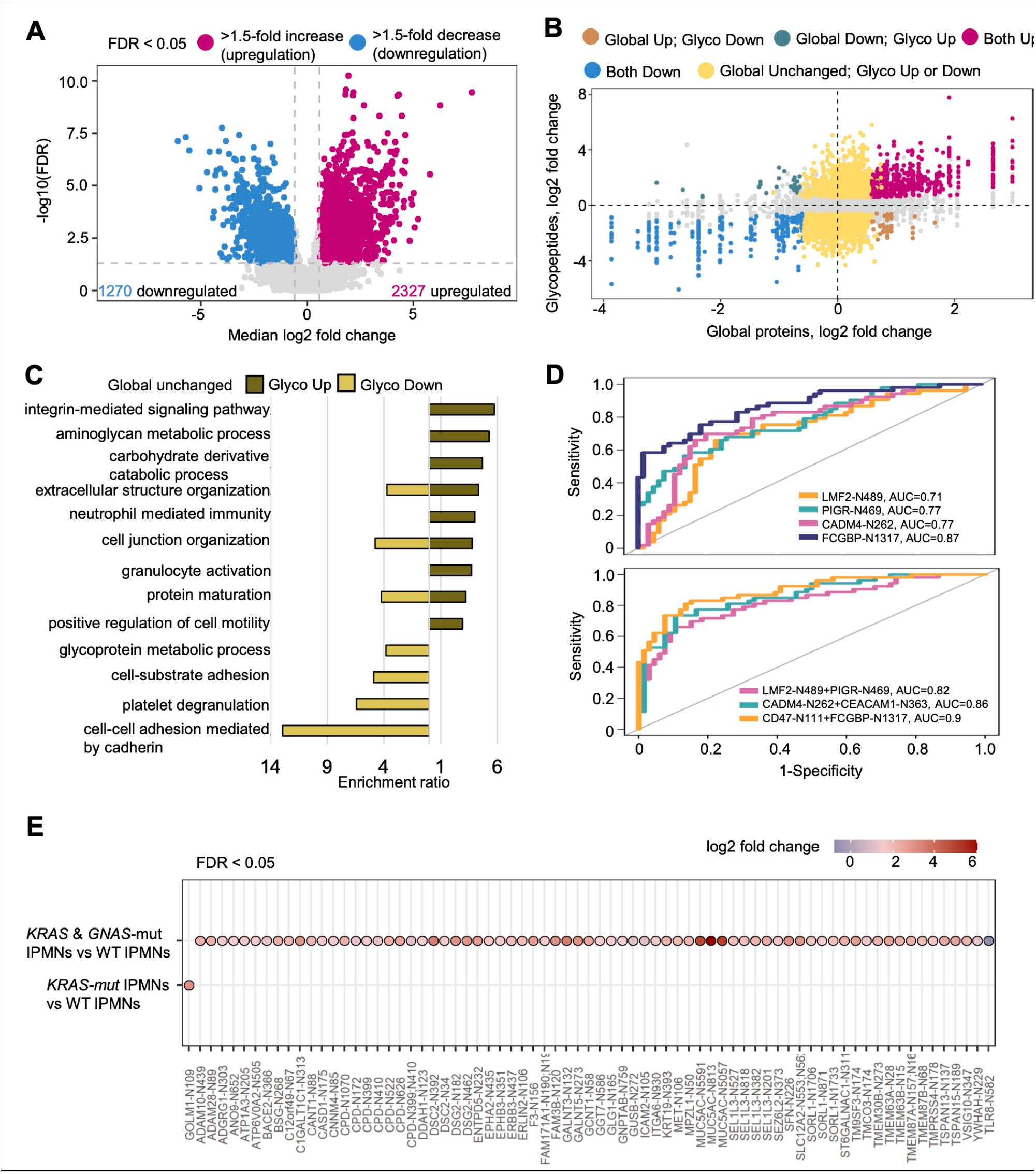
Altered glycosylation in 53 IPMNs. **A**. Comparison of glycopeptides in 53 IPMNs and 66 NDs. **B**. Expression changes in glycopeptides in comparison to the changes in global protein levels (IPMN vs NDs). **C**. Significantly enriched GO biological functions of upregulated and downregulated glycopeptides in instances in which the proteins were unchanged at the global level. **D**. Examples of individual glycopeptides and combination of glycopeptides capable of differentiating IPMNs from NDs. **E**. glycopeptides with differential expression in *KRAS*-mutant IPMNs or both *KRAS*- and *GNAS*-mutant IPMNs compared to *KRAS*-wildtype (WT) and/or *GNAS*-WT IPMNs.

Some cell surface glycoproteins, such as CEACAM5^66^, integrins (ITGA5, ITGB1)^67,68^, and mucins (including MUC1, MUC2, MUC4, MUC5AC, MUC5B, MUC6, and MUC13)^48,69^ that were overexpressed are known to facilitate cell adhesion and form protective barriers around the tumor surfaces. Additionally, CD276^70,71^, CD47^72,73^, and other immune-related glycoproteins were also upregulated in the IPMNs. This study is one of the first to report the upregulation of CD276 and CD47 in IPMNs, although they were previously discovered to be PDAC-associated proteins ^70-74^(Supplementary Table S3). Similar patterns of glycoprotein expression were observed when only the IPMNs with a somatic mutation were considered (Supplementary Fig. S3A and Table S3B). The glycoproteins expressed in the IOPNs are separately listed in Supplementary Table S3C.

More than 130 proteins were found to be upregulated at both the global protein and glycoprotein levels (Supplementary Tables S3D). These included mucins (MUC5AC, MUC6, and MUC13) and mucin-type O-glycan synthetic enzymes (C1GALT1C1, GALNT3, GALNT6, and GCNT1). More interestingly, extracellular matrix (ECM) interaction proteins^75^ such as CD47^73,74^, integrins (ITGA2, ITGA9, ITGB4, ITGB6, and ITGB8)^76^, and laminins (LAMA3, LAMB3, and LAMC2)^77^, were also upregulated at both the global protein and glycoprotein levels, strongly indicating changes in the tumor microenvironment in IPMNs.

Comparison of glycopeptide expression to global protein expression revealed that some changes in expression were only observed at the glycopeptide level, while others were only observed at the protein level (Fig. 3B, Supplementary S3D). This finding highlights the importance of characterizing both proteins and glycopeptides, as protein expression may not always equate to glycosylation modification^78^. To reveal the unique pattern of glycoprotein expression compared to protein expression in IPMNs, we analyzed the biological function (GO terms) of upregulated or downregulated glycopeptides whose global protein expression remained unchanged (Supplementary Table S3E). Some upregulated glycoproteins functioned in integrin-mediated signaling and leukocyte migration, indicating changes in the tumor microenvironment (Fig. 3C). Metabolic-related functions were also upregulated in the IPMN glycoproteome in instances where the global protein expression remained unchanged. Downregulated glycoproteins included those involved in cell-cell adhesion mediated by cadherin, an important modulator of epithelial-mesenchymal transition (EMT) during tumor progression^79^.

A comparison of glycopeptide expression in 53 IPMNs to 69 NDs revealed a number of glycoprotein changes associated with IPMNs (Supplementary Fig. S3B, Supplementary Table S3F). In particular, several glycopeptides and combinations of glycopeptides produced a mean bootstrap area under the curve (AUC) > 0.7 (Supplementary Fig. S3B and Table S3F-S2G) in distinguishing IPMNs from NDs. Including upregulated and downregulated glycopeptides, individual markers, including CADM4-N262, FCGBP-N1317, LMF2-N489 and PIGR-N469, achieved AUCs between 0.71 and 0.87 (Fig. 3D). These AUCs improved to as high as 0.9 when several markers were combined (Fig. 3D).

We next compared the patterns of glycopeptide expression in 17 IPMNs that harbored mutations in *KRAS* and *GNAS* to the 15 wildtype IPMNs (the term “wildtype IPMNs” in following content and figures refers to IPMNs that lacked any somatic mutations) (Fig. 3E). There were 71 glycopeptides involving 56 glycoproteins upregulated in IPMNS with both KRAS and GNAS mutations compared to wildtype IPMNs (Supplementary Table S3H-S3I). Among these 56 glycoproteins, O-glycosylation enzymes such as C1GALT1C1, GALNT3, GALNT5, GCNT1, and ST6GALNAC1 were overexpressed. This pattern could be explained by the activation of the PI3K-AKT signaling pathway in the cells with mutations. PI3K-AKT pathway-related glycoproteins, including EPHA2, ERBB3, ITGA6, MET, and YWHAH, were also observed among these 56 upregulated glycoproteins, as were transporter glycoproteins (ATP1A3, ATP6V0A2, CNNM4) related to inorganic ions, such as Na^+^/K^+^, H^+^, and Mg^2+^. The simultaneous upregulation of these ion transporter proteins suggests that the neoplastic cells in IPMNs with mutations in KRAS or GNAS may undergo significant adjustments in ion balance and may reprogram metabolic processes to meet high metabolic demand.

### Dysregulated carboxypeptidases associated with elevated levels of non-tryptic peptides

One function of the pancreas is to aid in digestion, and the normal pancreas produces digestive enzymes, including lipases, proteases, and amylases. The proteases secreted by the pancreas include trypsin, chymotrypsin, and carboxypeptidase. To investigate the protease expression levels in IPMNs, we found that the abundance of CPA1 and CPA2, the major carboxypeptidases in the pancreas, was higher in NDs compared to IPMNs, as were the levels of the chymotrypsins CTRL and CTRC (Fig. 4A).

**Figure 4.**
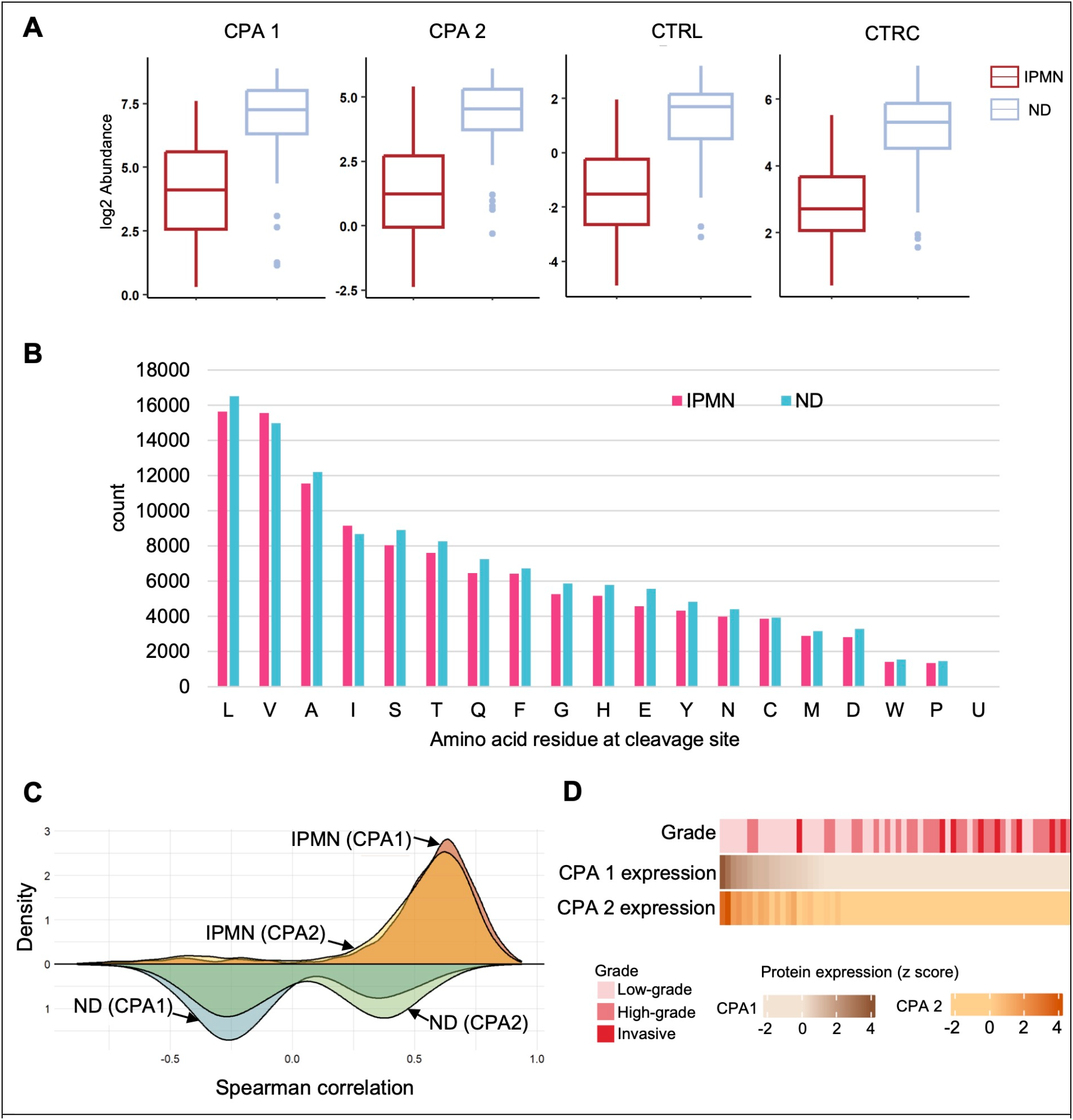
Carboxypeptidase (CPA1 and CPA2) and chymotrypsin (CTRL and CTRC) in association with IPMN and normal duct (ND) tissues. **A**. Expression profile of CPA1, CPA2, CTRL, and CTRC in IPMNs and NDs. **B**. Number of peptides with non-tryptic (K/R) ends in IPMNs and NDs. **C**. Distribution of spearman correlation between non-tryptic peptides and CPA1, and CPA2 in IPMNs and NDs. **D**. Protein expression of CPA1 and CPA2 in relation to IPMN grade.

These proteases are normally inactive zymogens in the pancreas and are activated in the duodenum by enteropeptidases^80^. However, prematurely activated proteases can generate peptides that can be detected by our mass spectrometry analysis. Due to the use of trypsin in our proteomic procedure, the tryptic peptides generated by endogenous trypsin could have been masked by the exogenous trypsin added to the samples. In contrast, chymotrypsin and carboxypeptidases can create non-tryptic peptides in addition to tryptic peptides. Herein, we discovered that non-tryptic peptides were present at surprisingly high levels in IPMN and ND samples (Fig. 1E). Semi-tryptic searches revealed that nearly all N-termini of the digested peptides had lysine or arginine residues, indicating these non-tryptic peptides were primarily generated by carboxypeptidase rather than chymotrypsin. The C-termini of the digested peptides were abundant with leucine, valine, and alanine residues, suggesting dysregulated carboxypeptidase activity (Fig. 4B).

To further explore the differences in non-tryptic peptides between IPMNs and NDs, we compared all non-tryptic peptides that significantly overlapped between IPMNs and NDs (p < 0.05) and their association with CPA1 and CPA2 levels, based on linear regression analysis (Supplementary Table S4A-S4B). From the distribution of Spearman correlations (Figs. 4C), the non-tryptic peptides strongly correlated with carboxypeptidase levels in the IPMN samples.

However, these correlations in ND samples were weaker than those in IPMNs or they even became negative. Subsequently, we elucidated the expression profiles of CPA1 and CPA2 in high-grade compared to low-grade IPMNs (Fig. 4D). The IPMNs with low-grade dysplasia had high levels of carboxypeptidase expression. High-grade IPMNs tended to have low levels of carboxypeptidase expression. The altered levels of proteases CPA1 and CPA2 associated with IPMN grade can be determined at global enzymatic level. While abnormal enzyme activation may occur in pancreatitis, presumably including neoplasm-initiated pancreatitis, we cannot rule out the possibility that the presence of non-tryptic peptides may be an artifact of abnormal carboxypeptidase activation during warm ischemia ^81,82^.

### Changes in protein expression patterns during neoplastic progression

We selected 104 of the most cellular (tumor cellularity >15%) PDACs and 43 their matched NATs from a previously reported series of PDACs and reanalyzed them using DIA-MS ^33^ (Fig. 5). In principal component analysis (PCA), the PDACs, IPMNs, and normal samples (NDs and NATs) clustered separately (Fig. 5A). Furthermore, the IPMN samples clustered between PDACs and normal samples, supporting the concept that IPMNs represent a transitional stage from normal ductal tissue to invasive cancer^83,84^. Comparing the protein expression in the 104 PDACs to the matched 43 NATs revealed 1,539 proteins significantly upregulated more than 1.5 fold, and 902 proteins significantly downregulated more than 1.5 fold in the cancers (Fig. 5B, Supplementary Table S5A). The vast majority of these proteins have been reported previously, and a number of them, such as S100P, THBS2, and SERPINB5 were also identified in this study as overexpressed in IPMNs (Figs. 2A and 5B). To determine the compartment (neoplastic cell vs. stroma) in which these proteins were expressed, we again utilized a previously published database (Supplementary Table S2I)^54^. Similarly to what was done with the IPMNs, we also validated the expression of select proteins (S100P, SERPINB5, and THBS2) and the compartment in which these proteins were expressed in PDAC tissues using immunohistochemistry (Fig. 5C, Supplementary Table S2J)^52,53^. In specific, we found that S100P and SERPINB5 were expressed by neoplastic invasive cancer cells, while THBS2 was expressed by stromal cells.

**Figure 5.**
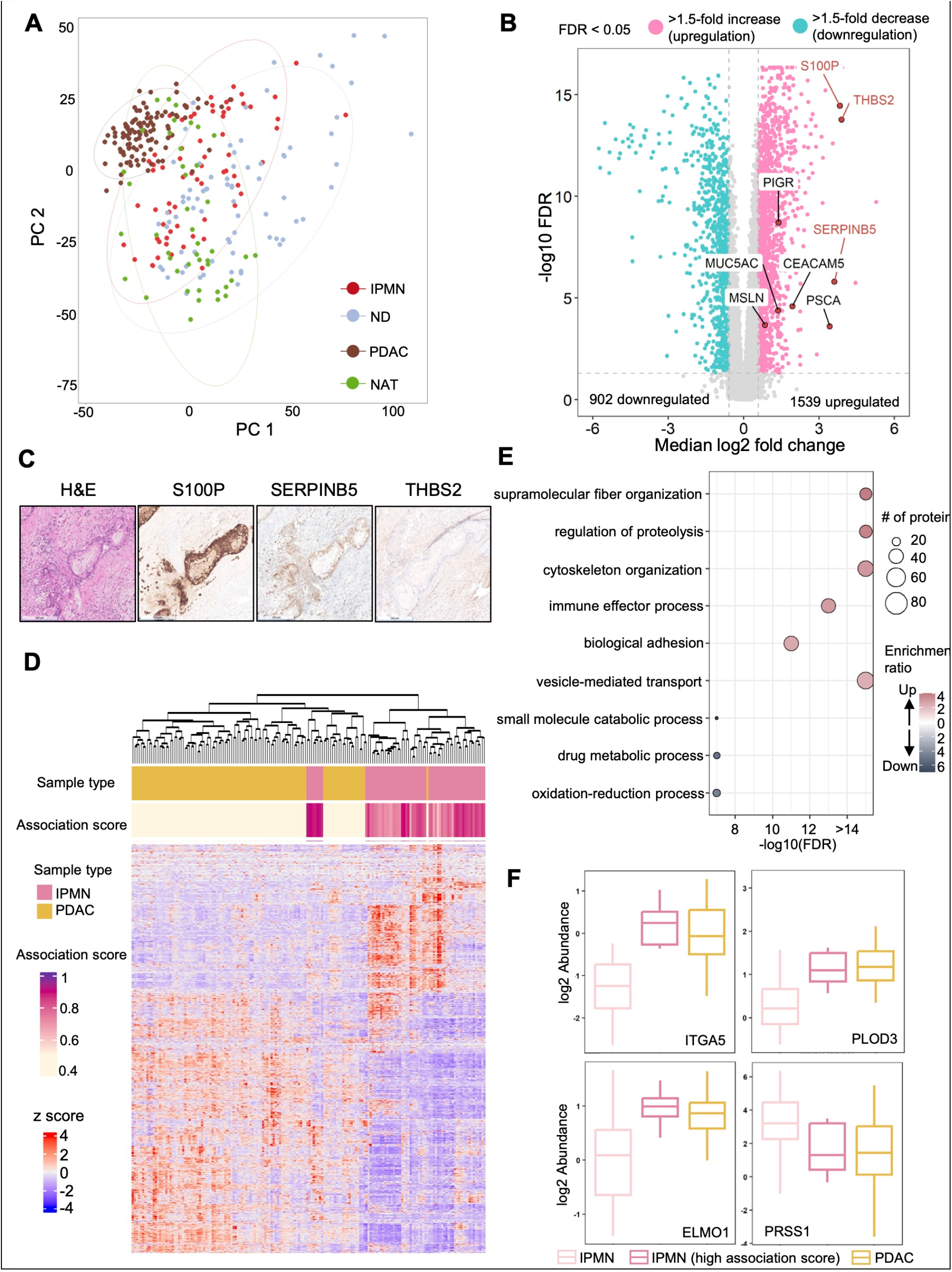
Proteomic comparison between IPMNs and PDACs with high tumor cellularity. **A**. PCA analysis of the cohort. **B**. Differentially expressed proteins in 104 PDACs relative to 43 NATs. Proteins with immunochemistry labeling are in red. Other labeled proteins also showed upregulation in IPMNs vs NDs in Figure 2A. **C**. IHC labeling results of S100P, SERPINB5, and THBS2 in PDAC tissue. **D**. Hierarchical clustering of PDACs and IPMNs, where 8 IPMNs with high association scores group with PDACs. **E**. Enriched GO biological processes using overexpressed and downregulated proteins in PDACs and IPMNs grouped with PDACs compared to the remaining IPMNs. **F**. Examples of the proteins showing significant changes between PDACs and IPMNs grouped with PDACs relative to other IPMNs.

To further investigate which IPMNs are more associated with PDACs, we first identified 1,670 proteins significantly changed in PDACs compared to NATs and normal ducts and calculated an association score between PDACs and IPMNs based on these proteins (Supplementary Table S5B-S5C). We found that 8 IPMNs clustered with PDACs (Fig. 5D), each having a relatively high association score (≥ 0.88). We further examined the 1,670 proteins and found 351 proteins that were overexpressed in PDACs and in some IPMNs that grouped with PDACs compared to the remaining IPMNs (Supplementary Table S5D). These proteins are potential markers of IPMN progression since they were elevated in IPMNs with higher PDAC association scores. Among these 351 proteins, those related to invasion and metastasis, such as cell adhesion, cytoskeletal dynamics, and cell motility, warrant more investigation (Fig. 5E and Supplementary Table S5E). For instance, the expression of ITGA5, PLOD3, and ELMO1 was higher in IPMNs with high association scores and in PDACs than in IPMNs without high association scores (Fig. 5F)^63,68,85^. On the other hand, we found 106 proteins that were downregulated in PDACs and IPMNs with high association scores relative to remaining IPMNs, and these were enriched in metabolic processes (Fig. 5E). As expected, PRSS1 expression decreased in IPMNs with high association scores compared to those without high association scores, consistent with its function as a normal digestive enzyme (Fig. 5F). These 351 upregulated and 106 down regulated proteins are potential markers for the early detection of IPMN with high progression potential.

### Proteomic subtyping highlights the heterogeneity of intraductal papillary mucinous neoplasms

Based on the morphologic features of the neoplastic epithelium, IPMNs can be classified into three subtypes: gastric-type, intestinal-type, and pancreatobiliary-type^86,87^. Furthermore, IPMNs can be categorized as having low-grade dysplasia, high-grade dysplasia, and as having an associated invasive cancer, based on the degree of cytoarchitectural atypia and the presence or absence of an associated invasive cancer^86,87^. However, these subtypes or classifications are not based on molecular characterization. Herein, we applied proteomics based non-negative matrix factorization (NMF) subtyping strategies to the IPMNs and PDACs to explore tumor heterogeneities and reveal progression features^88^. This analysis showed three clusters (Fig. 6A and Supplementary Table S6A-S6B), NMF 1 was entirely composed of PDACs. NMF 2 included some PDACs and some IPMNs, while NMF 3 was almost entirely composed of IPMNs with only one PDAC. Most (75%) IPMNs with an associated PDAC were clustered in NMF 2. IPMNs without KRAS or GNAS mutations also tended to cluster in NMF 2. KEGG pathway enrichment using signature proteins distinguished these three clusters (Fig. 6B and Supplementary Table S6C). Notably, NMF 3 has its own features that enriched in amino sugar metabolism, metabolic processes, and pancreatic secretion. Several mucin glycosylation related enzymes in NMF 3, such as UDP-galactose-4-epimerase (GALE), Glutamine-fructose-6-phosphate aminotransferase 1 (GFPT1), GNE, and Glucosamine-phosphate N-acetyltransferase 1 (GNPNAT1), are involved in amino sugar metabolism. The sodium/potassium-transporting ATPase (ATP1A1, ATP1B1) and ADP/ATP translocase (SLC25A4) found in NMF 3 are key players in metabolic processes. Ras-related proteins (RAB27B, RAB3D) found in NMF 3, which control the maturation, trafficking, and exocytosis of secretory vesicles are highlighted in pancreatic secretion. These unique features of NMF 3 indicate reprogrammed metabolic processes associated with increased mucin glycan synthesis and extracellular vesicle secretion.

**Figure 6.**
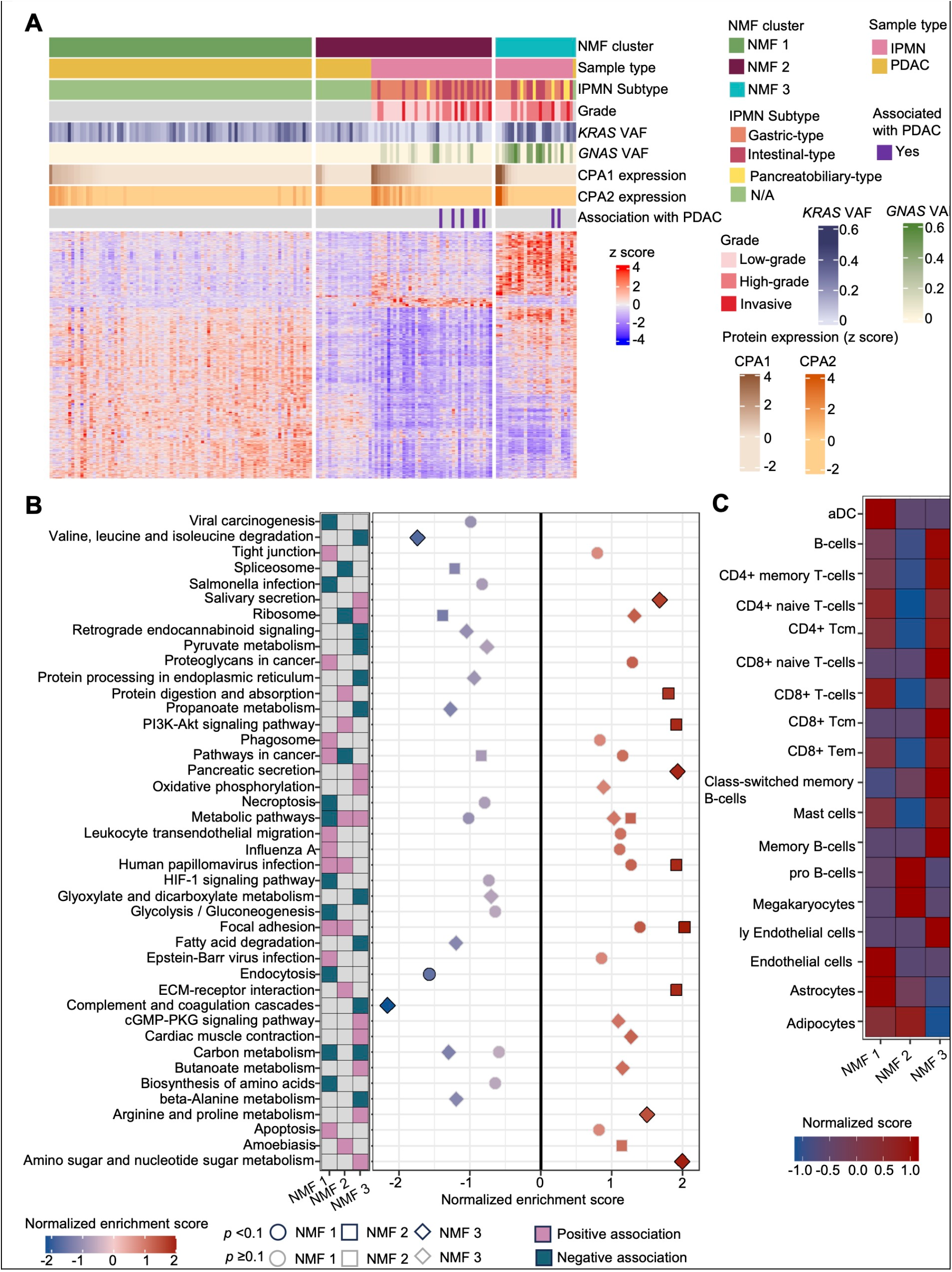
Proteomic subtyping of IPMNs and PDACs with high tumor cellularity. **A**. Three subtypes derived via NMF clustering. **B**. KEGG pathway enriched by using signature proteins differentially expressed in one NMF cluster compared to the remaining NMF clusters. **C**. Cell type enrichment based on the gene signatures from xCell for each NMF cluster.

Major differences between NMF 2 and NMF 3 include elevated collagen and matrix proteins (COL4A1, COL4A2, COL4A5, COL6A1, COL6A2, COL6A3, TNXB), which are enriched in the pathways of ECM-receptor interaction, PI3K-Akt signaling, and others (Fig. 6B). These findings highlight the reorganization of the ECM interactions during the progression from IPMNs to PDACs. Gelatinase A (MMP2) and Fibronectin (FN1), along with actin (ACTB, ACTN1) and the cytoskeleton protein (MYH9), profile the cell adhesion and migration in pathways of cancer and proteoglycans in cancer. These associations are not shown or are negatively associated in NMF 3. Our molecular level clustering analysis identifies some of the differences between the molecular features of IPMNs and those of PDACs, providing new insights towards the progression from non-invasive precursor lesions to invasive pancreatic cancer.

### Intraductal papillary mucinous neoplasms immune protein signatures

During progression from non-invasive lesions to invasive cancer, protein patterns related to the tumor microenvironment change significantly. To better understand the dynamics and features of the microenvironment in different NMF clusters, we classified microenvironment protein signatures, particularly focusing on immune protein signatures and the degree of immune infiltration (Fig. 6C)^89^. This approach aimed to highlight the potential for targeting immune modulators in the treatment of high-grade or invasive IPMNs. We annotated the samples in the NMF 3 as “immune hot” neoplasms due to the infiltration of various CD8+ T cell subtypes. The NMF 1 cluster, entirely composed of PDACs, exhibits “cold” signatures with CD8+ native T-cells and CD8+ central memory T cells (CD8+ Tcm), which suggests an exhausted immune response and could lead to immune evasion by the invasive cancer^90,91^. These immune protein signatures add additional value to the molecular level clustering, providing novel insights for understanding cancer progression.

## Discussion

Despite the introduction of immune and other targeted therapies^92^, invasive pancreatic cancer continues to have an extremely high mortality rate ^1,93^. Although the detection and surgical resection of non-invasive precursor lesions, including intraductal papillary mucinous neoplasms, intraductal tubulopapillary neoplasms, and intraductal oncocytic papillary neoplasms (here collectively referred to as IPNs), offers the hope of curing a neoplasm before it has the opportunity to progress to incurable disease, the surgical resection of precursor lesions risks overtreatment ^18,19^. Today, even with advances in imaging and new tests to detect tumor-derived somatic mutations, many lesions surgically resected because they are clinically believed to be intraductal papillary mucinous neoplasms with high-grade dysplasia turn out to be other types of neoplasms or IPMNs with only low-grade dysplasia ^18,19,94^.

To understand the drivers of IPNs and identify novel markers, we characterized these neoplasms in-depth using state-of-the-art mass spectrometry to achieve deep proteome and glycoproteome coverage. To ensure unbiased profiling, we selected IPNs with different genomic backgrounds, grades of dysplasia, and histologic subtypes (Fig. 1), and compared the expression patterns in these neoplasms with those in macrodissected normal ducts. We focused primarily on IPMNs as they are the predominant type of IPNs. A number of proteins and glycoproteins were identified that are expressed in higher levels in IPMNs than in normal pancreatic ducts (Figs. 2A and 3A, and Supplementary Tables S2A and S3A). As expected from a tumor with “mucinous” in its name, these included proteins involved in glycan biosynthesis as well as a number of known surface glycoproteins such as CEACAM5, MUC1, MUC2, MUC4, MUC5AC, MUC6, and MUC13^69,83^. These findings, coupled with the validation of selected markers by immunolabeling, help corroborate our approach. The various overexpressed proteins and glycoproteins identified are potential biomarkers for this tumor type as many can be detected in clinically obtainable cyst fluid (Figs. 2E and 3D).

Of interest, unlike mass spectrometry analyses of other tumor types, we identified a large number of non-tryptic peptides (Fig. 1E and 4). These highlight the dysregulation of proteases, particularly carboxypeptidase in our samples. These non-tryptic peptides were most abundant in normal duct and IPMNs with low-grade dysplasia samples. Presumably the high-grade IPMNs, as they are more likely to be mass-forming lesions that protrude into the pancreatic ducts, are more likely to block the flow of digestive enzymes^95,96^. The level of carboxypeptidases may be clinically used to distinguish between the IPMN subtypes (Fig 4D).

To investigate which IPMN-elevated proteins were associated with PDAC, we re-analyzed a previously well-characterized set of PDACs and NATs using the same mass spectrometry technology used in analyzing the IPNs, NDs and cyst fluid samples in this study. This provided a unique opportunity to define the changes in expression that occur throughout the entire progression from normal duct, through precursor lesion, to invasive cancer^33^. We found that the normal samples (NDs and NATs), IPMNs, and PDACs, clustered separately in principal component analysis (PCA) (Fig. 5A), and, as expected, the IPMN samples clustered between PDACs and normal samples. This finding supports the hypothesis that IPMNs represent a transitional stage from normal ductal tissue to invasive cancer^83,84,97,98^.

Furthermore, we identified 1,670 proteins that significantly changed in PDACs compared to normal samples. Using these 1,670 proteins, we calculated an association score between PDACs and IPMNs, and found 8 IPMNs clustered with PDACs giving high association scores. Among these, 351 proteins were overexpressed in PDACs and the 8 IPMN that clustered PDACs, while 106 proteins were downregulated; these proteins are potential markers for early detection and understanding the progression of IPMNs with high PDAC association scores, and as noted earlier, many can be detected in clinically obtainable cyst fluid (Fig. 2E).

Two methods, mining a previously reported database and immunolabeling, were employed in this study to classify the cellular compartment (neoplastic cells or stroma) in which the various proteins and glycoproteins were likely expressed (Figs. 2C and 5C, Supplementary Tables S2J)^99,100^. Defining cellular compartments not only helps clarify function of these proteins and glycoproteins, but also indicates which proteins and glycoproteins (those expressed by the neoplastic cells) are more likely to be clinically detectable in cyst fluid samples, making them more clinically useful as markers.

To investigate the molecular level progression of invasive cancer, we conducted proteomics based NMF clustering within IPMNs and PDACs. We identified three clusters, NMF 1 (PDACs), NMF 2 (PDACs and IPMNs), and NMF 3 (mostly IPMNs) (Fig. 6A). The IPMNs with high association scores with PDACs (Fig. 5D) are also most likely grouped with PDACs in NMF 2, which validates our previous analysis. NMF 3 was enriched in metabolic processes, N-glycosylation processes and pancreatic secretion. NMF 2 suggested the re-shaping process of ECM, while NMF 1 showed invasive features, highlighting the progression from IPMNs to PDACs. NMF cluster based immune cell type signatures provide a window for potential immunotherapy in high-grade IPMN patients (Fig. 6C). This molecular level clustering analysis provides an opportunity to observe the heterogeneity among different IPMN subtypes, offering insights into the progression processes. In terms of glycosylation, the mucin-type O-glycan synthetic enzymes (C1GALT1C1, GALNT3, GALNT6, and GCNT1) are elevated from NDs to IPMNs, while other O-glycan synthetic enzymes (PLOD3, GXYLT1, UAP1, GNE, and B3GNT3) increase in high-grade IPMNs. The transition from IPMNs to PDACs can be distinguished by N-glycosylation enzymes (GFPT1, GNE, GNPNAT1, PMM2, and UAP1). Regarding ECM changes, ECM interaction proteins (CD47, integrins, and laminins) are upregulated from NDs to IPMNs, and ECM proteins such as collagen proteins and matrix proteins are elevated from IPMNs to PDACs. Proteins related to cancer invasion and metastasis (ITGA5, PLOD3, ELMO1, MMP2, and FN1) are overexpressed as the disease progresses from low-grade IPMNs to high grade-IPMNs and ultimately to PDACs.

We also found that IPMNs with both *KRAS* and *GNAS* somatic mutations exhibit particularly elevated glycoprotein processing proteins, along with activation of the PI3K-AKT signaling pathway^101^. These changes may lead to significant adjustments in ion balance and metabolic processes suggested by the simultaneous increase in ion transporter proteins such as ATP1A3 and CNNM4.

There are several potential weaknesses of this study that should be acknowledged. We relied on surgically resected tumors, and many of the neoplasms included in this study were low-grade tumors. We therefore paired high-grade IPMNs with IPMNs with an associated invasive cancer in this study. Further studies focusing on purely high-grade IPMNs and, in particular, on the changes in expression that occur early in the transition from high-grade dysplasia to invasive carcinoma are warranted. In addition, the prevalence of somatic mutations identified in our IPNs was slightly lower than anticipated from the literature, perhaps reflecting the fraction of neoplasms with low-grade dysplasia. We believe that this didn’t bias our results, as sub-analyses of the expression patterns when only the neoplasms with somatic mutations were included yielded similar results as when all neoplasms were included (Supplementary Figs. S2D and S3A).

In summary, we present an in-depth characterization of the patterns of protein and glycoprotein expression during the progression from normal duct to intraductal papillary neoplasm, to invasive pancreatic cancer. This understanding forms the basis for evidence-based approaches to early detection and novel treatment strategies. We envision that analyzing a panel of somatic mutations and protein/glycoprotein markers in cyst fluid will improve the management of patients with pancreatic cysts^28^.

## Methods

### Subject details

Fresh tissue, including 76 macro-dissected normal main pancreatic ducts and 69 intraductal papillary neoplasms (64 IPMNs and 5 IOPNs), 5 serous cystic neoplasms (SCNs), 4 mucinous cystic neoplasms (MCNs), and 32 cyst fluid samples (22 matched with analyzed IPMNs), was harvested from 126 patients (59 males, 67 females, age range 21 to 87) from five different countries who underwent surgical resection for a pancreatic lesion. Cyst fluid was aspirated at the surgical pathology bench from 22 of the analyzed IPMN-matched patients, 2 were from other IPMN patients, 3 IOPN patients, and 5 other IPMN cyst samples. The other 5 cyst fluid samples were from other pancreatic neoplasms including 2 MCNs, 2 SCNs and 1 solid pseudopapillary neoplasm. Of the 76 main pancreatic duct specimens, 32 were harvested from the 64 patients with an IPMN, and 44 were harvested from patients who underwent surgery for a neoplasm that does not involve the duct system. In addition, 104 PDAC fresh-frozen samples with >15% neoplastic cellularity and 43 PDAC matched NATs were obtained from a previous multi-omics study of 144 PDACs cohort and rerun on the same platform as the normal duct, IPMN and cyst fluid samples (DIA-MS technology) ^33^. The hematoxylin and eosin-stained sections of all IPMNs were reviewed by one pathologist (T.F.) and the diagnosis, direction of differentiation, and histologic grade confirmed. 15 independent formalin-fixed and paraffin-embedded IPMNs were selected from the files of the Johns Hopkins Hospital for immunolabeling (Supplementary Table S2J).

### Sample processing for protein extraction from tissues, tryptic digestion, and global proteomic sample preparation

All tissue samples for this study were processed for mass spectrometry (MS) analysis at Johns Hopkins University. The sample preparation for global proteomic analysis followed the previous described approach in PDAC study, and glycoproteomic analysis was carried out by liquid handing systems as previous published work^33^. All tumor and normal ductal tissue samples were cryo-pulverized and lysed in urea lysis buffer as previously published standard protocol ^102^ (8 M urea, 75 mM NaCl, 50 mM Tris pH 8.0, 1 mM EDTA, 2 µg/mL aprotinin, 10 µg/mL leupeptin, 1 mM PMSF, 10 mM NaF, phosphatase inhibitor cocktail 2 [1:100 dilution], phosphatase inhibitor cocktail 3 [1:100 dilution], and 20 µM PUGNAc) by sonicating for 30s x 3 in ice water). Tissue debris was removed by centrifugation at 16,000 x g for 15 min at 4 °C. The lyzed protein supernatant was collected, the concentration was measured by BCA assay and adjusted to 8 mg/mL in 8M urea lysis buffer. The lyzed proteins were reduced with dithiothreitol (6 mM, 37 °C, 1h) and alkylated by iodoacetamide (12 mM, room temperature in dark, 45 min). After that, the reduced proteins were diluted to 2M urea concentration with 50 mM Tris-HCl pH 8.0 buffer, digested by Lys-C (enzyme to protein ratio is 1 mAU:50 mg, 2h at room temperature) and trypsin (enzyme to protein ratio is 1 mg:50 mg, overnight incubation at room temperature), respectively. The proteolytic reactions were quenched by 50% formic acid to adjust pH < 3. About 5 µg digested peptides were aliquoted for global proteomics analysis by stage tip desalting and dried using Speed-Vac (Thermo Scientific). 1 µg peptides were loaded on Evotip (Evosep Biosystems) according to the EvosepOne (Evosep Biosystems) protocol for global proteomic analysis on timsTOF-HT (Bruker). The remaining digested peptides were desalted on reverse phase C18 96-well plate and dried using Speed-Vac for glycopeptide enrichment.

### Enrichment of intact glycopeptides by liquid handing systems from global peptides

As previously reported, C18/MAX-Tips were conditioned using acetonitrile, 100 mM triethylammonium acetate, 95% acetonitrile in 1% TFA, and 0.1% TFA, respectively in Versette Liquid Handling System (Thermo Scientific). The remaining digested peptides were reconstituted in 300 µL 0.1% TFA (trifluoroacetic acid), about 200 µg global peptides bound onto the C18/MAX-Tips (20 cycles) and washed with 0.1% TFA (10 cycles). After that, the non-glycopeptides were eluted from the C18/MAX-Tips with 200 µL 95% acetonitrile in 1% TFA (3 × 10 cycles), and glycopeptides were sequentially eluted with 200 µL 50% acetonitrile in 1% TFA (3 × 10 cycles). Samples were dried using Speed-Vac. The dried glycopeptide samples were resuspended in 40 µL 0.1% TFA and aliquoted 20 µL glycopeptide out into a 96-well plate. PNGaseF digestion buffer (2µL 1000 units PNGaseF added to 98 µL 100mM triethylammonium bicarbonate) was added to the 96-well plate and reacted at 37 °C overnight. The reaction was quenched by formic acid to adjust pH < 3 and cleaned up by stage-tip. The samples were dried and prepared for five injections and loaded one injection on Evotip (Evosep Biosystems) for glyco proteomic analysis on timsTOF-HT (Bruker).

### EVOSEP-TIMSTOF for global and glycoroteomic analysis

EvosepOne (Evosep Biosystems) LC system was coupled with timsTOF-HT mass spectrometry. Global peptides and glycopeptides were loaded on Evotip and separated on PepSep C18 column of 15 cm x 150 µm, 1.5 µm (Bruker) in Bruker column toaster (50 °C) at a 30 SPD gradient. The global proteomic data was acquired using the DIA-PASEF mode on timsTOF-HT with settings as follows: MS1 scan range of 100-1700 m/z; MS2 scan range of 338-1338 m/z, mass width 25.0 Da without mass overlap, 1 mobility window, 1/K0 range of 0.70-1.45 V.s/cm^2^, ramp time 85.0 ms. The glycoproteomic data was acquired using the DIA-PASEF mode on timsTOF-HT with settings as follows: MS1 scan range of 100-1700 m/z; MS2 scan range of 338-1488 m/z, mass width 25.0 Da without mass overlap, 1 mobility window, 1/K0 range of 0.70-1.45 V.s/cm^2^, ramp time 85.0 ms. The samples were randomized and blinded to the individual generating the data. Replications of pooled samples were used as quality control throughout the entire data generation process.

### Data Processing for Data Independent Acquisition Analysis

Spectral library was generated in Spectronaut^®^ 18.4 (Biognosys AG) by combination of all search archives from semi-specific and full-specific tryptic searches, which contained entire patient sample cohort. The Pulsar search settings for the global proteomic search archive generation were selected semi-specific (or full-specific) tryptic digestion with up to two missed cleavages within the length range of 7–52 amino acids. The fixed modification was on carbamidomethylation of cysteine (C). The variable modifications were on oxidation of methionine (M) and acetylation of protein N-terminal. The library generation settings were followed BGS factory settings with all PSM FDR, peptide FDR and protein FDR below 0.01. For the glycosite containing peptide library generation, a variable modification from N to D was added and other settings were the same as global proteomic library generation.

For the global proteomic search, all DIA files including generated from both tumor and normal samples were loaded and searched under full and semi specific combined library with BGS default factory setting. The proteins were identified and quantified by stripped sequence, and data underwent cross-run normalization.

For the glycosite containing peptide search, all DIA files including generated from both tumor and normal were loaded together and searched under full and semi specific combined library with BGS default factory setting. The peptides were identified and quantified by modified sequence, without cross-run normalization.

### Differential analysis

The differential analysis was carried out by calculating the median log2 fold changes for the following: (i) IPMNs vs NDs, (ii) HG vs LG IPMNs, (iii) HG IPMNs vs NDs, (iv) KRAS- and/or GNAS-mutant IPMNs vs *KRAS*- and/or *GNAS*-WT IPMNs, (v) Gastric-type IPMNs vs remaining subtypes of IPMNs, (vi) Intestinal-type IPMNs vs remaining subtypes, (vii) PDACs vs NATs, (viii) PDACs vs NDs, (ix) IPMNs (with high association score) and PDACs vs remaining IPMNs, and (x) one NMF cluster vs another NMF cluster on global proteomic or glycoproteomic level when applicable. The p-values were computed using two-sided Wilcoxon rank sum test and adjusted (false discovery rate, FDR) using Benjamini-Hochberg method.

### Enrichment analysis for KEGG pathway and GO terms

KEGG pathways and GO enrichment analyses were performed using the over-representation analysis on the WebGestalt (version 2019) with default setting and top 10 enriched terms were exported^103^. Only significantly upregulated and/or downregulated proteins/glycopeptides were used in the analyses.

### Protein abundance ranking for tissues and cyst fluids

Significantly upregulated proteins in IPMNs (vs NDs) with less than 60% missing values across all the cyst fluid samples were used to investigate the relation between IPMN tissues and cyst fluids. The protein abundances were z-score transformed and then ranked within each sample.

Association score calculation to determine whether an IPMN was more associated with PDAC A total of 1,670 proteins were differentially expressed in PDACs compared to NDs and NATs, which were used for calculating the association between PDACs and IPMNs. In brief, median abundance was computed for each protein across 104 PDAC tissue samples. Association score was determined by calculating the Spearman correlation between abundances of the aforementioned 1,670 proteins in an IPMN tissue sample and the median abundances of those 1670 proteins in PDAC.

### Non-negative matrix factorization (NMF)-based clustering analysis

To further analyze the heterogeneity of IPMNs, we utilized NMF to perform proteomic clustering, including both IPMN and PDAC tissue samples. The clustering approach was conducted similar to our previous works^33,36^. Briefly, NMF was used to perform unsupervised clustering of tumor samples based on the abundances of proteins (only proteins with CVs in >10% quantile were used in the analysis). The feature matrix was scaled and standardized, thus all features were represented as z-scores. Since NMF requires a non-negative input matrix, the feature matrix was further converted as follows: (i) create one data matrix with all negative numbers zeroed, (ii) create another data matrix with all positive numbers zeroed and the signs of all negative numbers removed, and (iii) concatenate both matrices resulting in a data matrix with positive values and zeros only. The resulting matrix was then subjected to NMF analysis leveraging the NMF R-package^88^. To determine the optimal factorization rank k (i.e., number of clusters), a range of k from 2 to 5 was tested using default settings with 50 iterations. The optimal factorization rank k=3 (i.e., NMF 1, NMF 2, and NMF 3) was selected since the product of cophenetic correlation coefficient and dispersion coefficient of the consensus matrix was the maximum compared to other tested ks. The NMF analysis was repeated using 500 iterations for the optimal factorization rank. A list of representative features for each NMF cluster was derived that differential analysis was carried out using cluster-specific features by comparing one NMF cluster to the remaining NMF clusters as described in the Differential analysis section. Heatmap of NMF clusters was generated using ComplexHeatmap (version 2.4.3).

### Immune protein signatures of each NMF cluster

The cell type gene signatures were obtained from xCell and we extracted protein abundances of these gene signatures from our global proteomic data^89^. In this study, we focused on the cell types that were enriched in the immune-hot PDAC samples in our previous publication to examine differences among NMF clusters due to the degree of immune infiltration for the IPMN^33^. We first found the association between cell type protein signatures and each NMF cluster. Based on the differential abundance (i.e., log2 fold change) between a NMF cluster and the remaining NMF clusters, a total of 112 protein signatures were obtained. Among the 112 protein signatures, 33, 41, and 38 proteins were significantly changed in NMF 1, NMF 2, and NMF 3, respectively. The abundance of each protein signature in a NMF cluster was weighted and signed by multiplying the protein abundances to the aforementioned log2 fold change. For each cell type in each NMF cluster, a cell type association score was calculated by summing up the weighted and signed abundances of the protein signatures. The cell type associated scores for each cell type were z-score normalized (i.e., normalized scores) among the NMF cluster.Sample processing for somatic mutation analyses

### DNA extraction, library preparation, and targeted next-generation sequencing

DNA was extracted from cryo-pulverized fresh frozen tissue using the QIAamp DNA FFPE Tissue Kit (Qiagen, Hilden, Germany) and a modified protocol^104^. In brief, tissue was enzymatically digested overnight, mechanically sheared (M220 Focused-ultrasonicator, Covaris, Woburn, MA), and DNA extracted following the manufacturer’s instructions. DNA concentration and base-pair length were quantified using an electrophoresis device (TapeStation System, Agilent, Santa Clara, CA).

Library preparation was performed using the SureSelect XT HS2 DNA System (Agilent, Santa Clara, CA) as suggested, with the modification of using half the probe volume that is indicated in the original protocol for preparing the probe hybridization mix in some samples. A customized panel of 12 established pancreatic driver genes *(BRAF, CDKN2A, CTNNB1, GNAS, KLF4, KRAS, PIK3CA, PTEN, RNF43, SMAD4, TP53, VHL*) was used for capture (Supplementary Table S1C). In total, our gene panel contained 1,115 probes across 142 genomic regions (Supplementary Table S1D). Prepared libraries were stored at -20 °C until sequencing. Libraries were pooled and sequenced on a MiSeq instrument (Illumina, San Diego, CA) using the MiSeq Reagent Kit v2 and generating 2x 150 base-paired reads.

### Data processing and mutation call

Following removal of duplicate reads, sequence reads were aligned to human genome 38 (hg38; Human_GRCh38.p13_MajChr) using NextGENe (SoftGenetics, State College, PA). After alignment, putative somatic mutations were filtered based on the following criteria: 1) a total coverage of at least 6 distinct reads at the mutation position; 2) at least 4 distinct reads supporting the mutation; 3) mutation allele frequency (MAF) of >5% for single base substitutions (SBSs), >10% for indels, and >20% for homopolymer indels. Software-based mutation calls were inspected with NextGENe Viewer (SoftGenetics, State College, PA) and further processed using Excel (Microsoft, Redmond, WA). Putative somatic mutations in genes covered in our targeted panel listed above were selected and alterations designated by NextGENe as ‘noncoding’, ‘synonymous’, or ‘none’ removed.

With the goal to identify somatic mutations in our tumor samples, 17 unmatched normal ductal samples that lacked *KRAS* and/or *GNAS* somatic mutations served as normal tissue for reference. Putative somatic mutations identified in normal samples were classified as germline variants. Putative somatic mutations unique to the tumor samples were visually inspected using Integrative Genomics Viewer (IGV; Broad Institute, Cambridge, MA) and compared to the reference genome hg38 to remove sequencing artifacts. In addition, hotspots for *KRAS* (codons G12, G13, Q61) and *GNAS* (codon R201) were visually inspected in all samples using IGV, irrespective of software-based mutation call by NextGENe. After visual inspection putative somatic mutations in genes other than *KRAS* or *GNAS*, that were not present in the Genome Aggregation Database (gnomAD, Broad Institute, Cambridge, MA; datasets GRCh38 gnomAD v4.0.0; gnomAD v3.1.2; gnomAD v3.1.2 non-cancer) or present in with an allele frequency <0.1% in all populations, were classified as somatic mutations. Putative somatic mutations present in these databases in any population at an allele frequency >0.1% were classified as germline variants.

### Immunohistochemistry

FFPE tissue blocks containing human primary tumors were sectioned at 4 mm onto Superfrost Plus microscope slides (VWR International, catalog 48311-703). Automated immunohistochemistry was performed at the Oncology Tissue and Imaging Services Core of the Johns Hopkins University School of Medicine using a Ventana Discovery Ultra automated slide staining system (Roche). Reagents used for deparaffinization, heat-induced epitope retrieval, and chromogenic signal detection were obtained from Roche and used according to the manufacturer’s protocol. Heat induced epitope retrieval (HIER) was performed using Ventana Ultra CC1 buffer (Roche, catalog 6414575001) at 96°C for 64 minutes. Following HIER, primary antibodies were at incubated at 37°C for 60 minutes under the following conditions: anti-S100P antibody (Abcam, ab124743, 1:2000 dilution), anti-SERPINB5 antibody (Abcam, ab182785, 1:200 dilution), and anti-THBS2 antibody (Abcam, ab112543, 1:100 dilution). Primary antibodies were detected using an anti-rabbit HQ and anti-HQ HRP detection system (Roche, catalog 7017812001 and 7017936001) followed by Chromomap DAB IHC detection (Roche, catalog 5266645001) according to the manufacturer’s protocol.

### Analysis of location of proteins overexpressed by IPMNs

A list of genes that are enriched in PDAC cells compared to adjacent stromal cells was obtained from a previously published study ^54^. Using this list, we searched for genes that corresponded proteins that were found to be upregulated in IPMNs compared to normal ducts and classified them as likely expressed in neoplastic cells or as likely expressed in stromal cells. Filtered results are summarized in Supplementary Table S2I.

## Acknowledgments

We thank the Dr. Masamichi Mizuma (Department of Surgery), Dr. Michiaki Unno (Department of Surgery), and Dr.Yuko Omori from (Department of Investigative Pathology) from Tohoku University Graduate School of Medicine for collecting tissues and clinical information. This study was generously funded by Susan Wojcicki and Dennis Troper. This work was also supported by the Clinical Proteomic Tumor Analysis Consortium (CPTAC) (U24CA210985 and U24CA271079) and Pancreatic Cancer Detection Consortium (U01CA274514) from the National Cancer Institute (NCI), National Institutes of Health (NIH).

## Supplementary Figures

**Figure S1.**
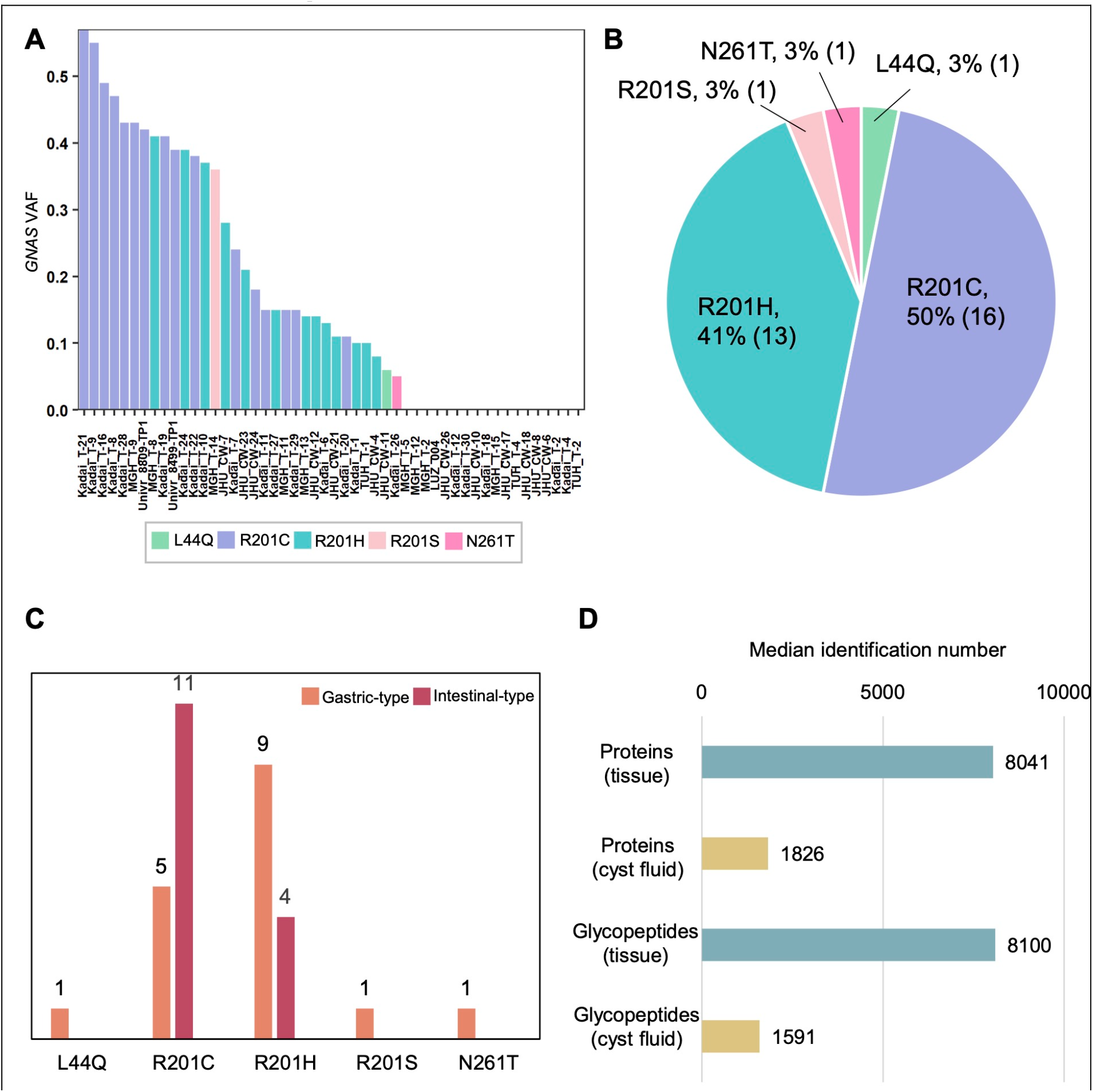
*GNAS* hotspot mutation in the IPNs. **A**. *GNAS* VAF distribution in 32 IPMNs. **B**. Distribution of *GNAS* hotspot mutations. **C**. Subtypes of IPMNs in association with each *GNAS* hotspot mutation. **D**. The median identification numbers in tissues and cyst fluids from global proteomics and glycoproteomics.

**Figure S2.**
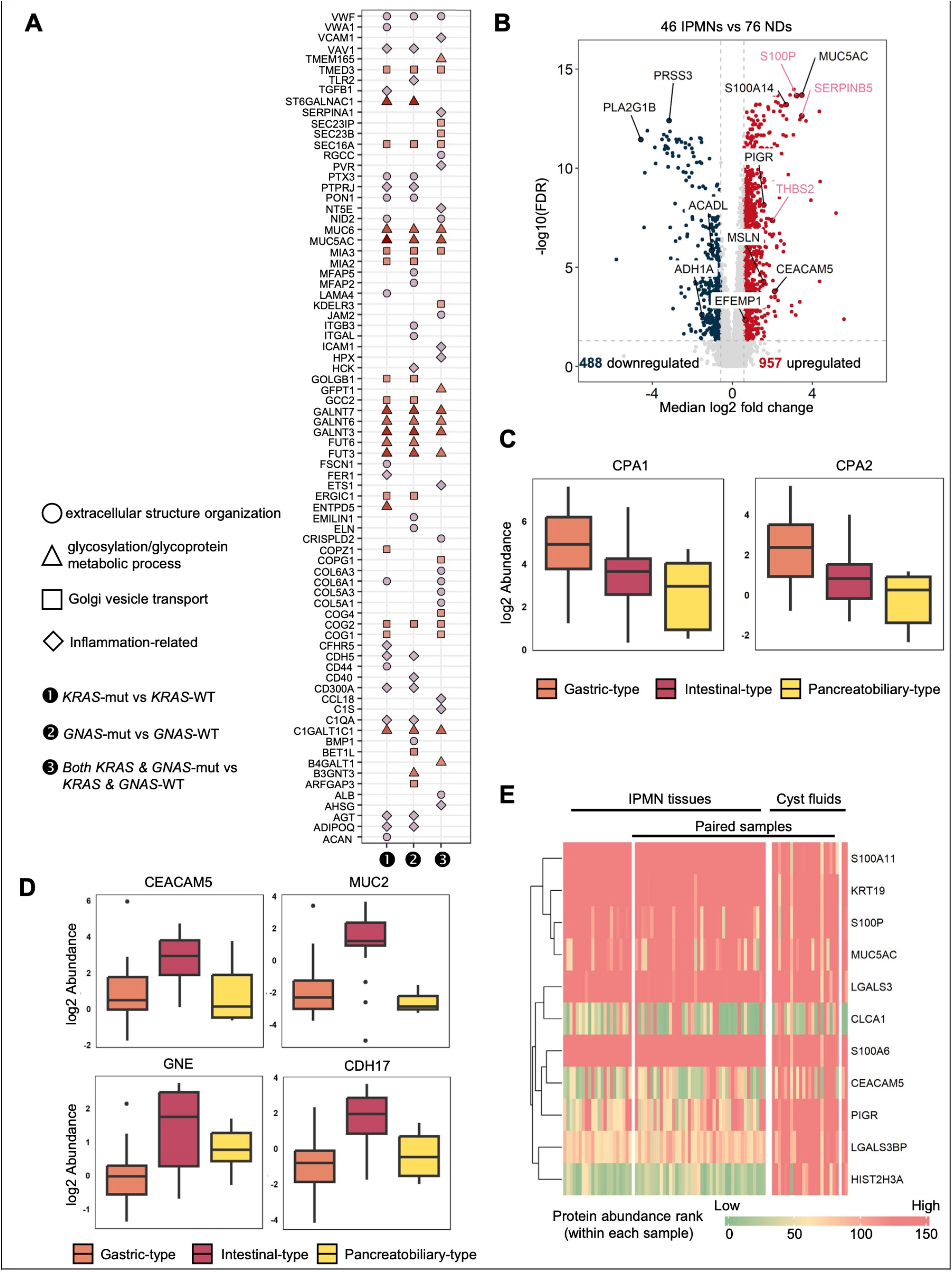
Potential of using cyst fluid to detect IPMNs and a comparison between *KRAS*/*GNAS*-mutant IPMNs and *KRAS*/*GNAS*-wildtype (WT) IPMNs. **A**. Top 10 proteins with significant fold changes in *KRAS*-mutant, *GNAS*-mutant, and/or both *KRAS*-& *GNAS*-mutant IPMNs vs *KRAS*/*GNAS*-WT IPMNs. These proteins were associated with biological processes of Golgi vesicle transport, glycosylation, inflammation, or extracellular structure organization. **B**. Comparison of 46 IPMNs (with at least one somatic mutation) and 76 NDs. Proteins with immunochemistry labeling are in pink. **C**. Examples of highly expressed proteins in gastric-type IPMNs compared to intestinal-type and pancreatobiliary-type IPMNs. **D**. Examples of highly expressed proteins in intestinal-type IPMNs compared to gastric-type and pancreatobiliary-type IPMNs. **E**. Expression profiles of proteins from Figure 2E in IPMNs and cyst fluids without paired samples in comparison to IPMNs with paired cyst fluids.

**Figure S3.**
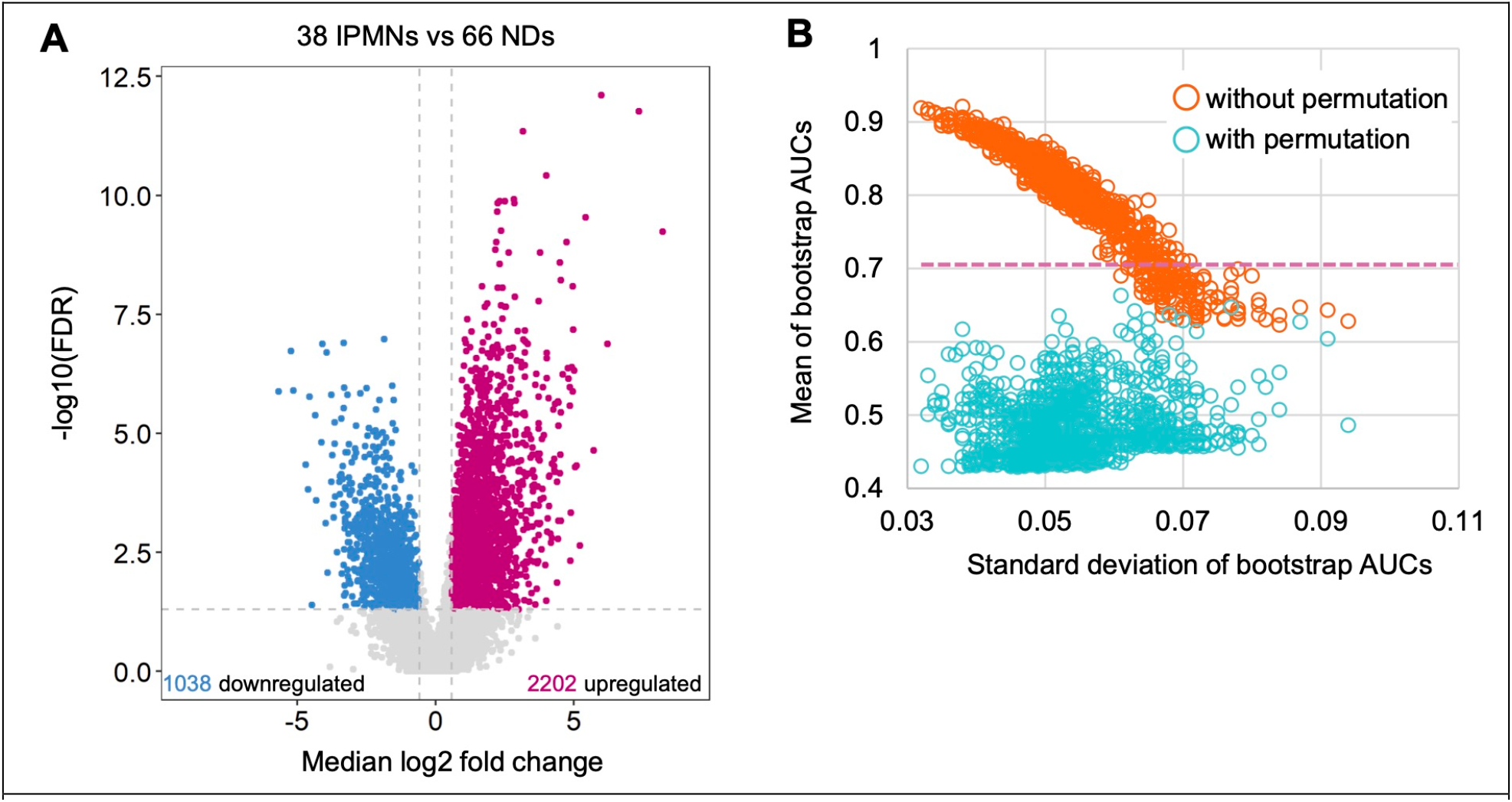
Glycopeptide expression changes in IPMNs with at least one somatic mutation and clinical utilities of glycopeptides. **A**. Differential analysis of 38 IPMNs and 66 NDs at glycopeptide level, where only IPMNs with at least one somatic mutation were considered. **B**. Individual and multi-glyco marker panels. Individual markers/panels with good performance compared to random models from label permutation are above the dashed line in pink.

## References

1. Siegel, R.L., Giaquinto, A.N., and Jemal, A. (2024). Cancer statistics, 2024. CA Cancer J Clin 74, 12–49. 10.3322/caac.21820.

2. Rahib, L., Smith, B.D., Aizenberg, R., Rosenzweig, A.B., Fleshman, J.M., and Matrisian, L.M. (2014). Projecting cancer incidence and deaths to 2030: the unexpected burden of thyroid, liver, and pancreas cancers in the United States. Cancer Res 74, 2913–2921. 10.1158/0008-5472.CAN-14-0155.

3. Rahib, L., Wehner, M.R., Matrisian, L.M., and Nead, K.T. (2021). Estimated Projection of US Cancer Incidence and Death to 2040. JAMA Netw Open 4, e214708. 10.1001/jamanetworkopen.2021.4708.

4. Quante, A.S., Ming, C., Rottmann, M., Engel, J., Boeck, S., Heinemann, V., Westphalen, C.B., and Strauch, K. (2016). Projections of cancer incidence and cancer-related deaths in Germany by 2020 and 2030. Cancer Med 5, 2649–2656. 10.1002/cam4.767.

5. Huang, J., Lok, V., Ngai, C.H., Zhang, L., Yuan, J., Lao, X.Q., Ng, K., Chong, C., Zheng, Z.J., and Wong, M.C.S. (2021). Worldwide Burden of, Risk Factors for, and Trends in Pancreatic Cancer. Gastroenterology 160, 744–754. 10.1053/j.gastro.2020.10.007.

6. Bretthauer, M., Loberg, M., Wieszczy, P., Kalager, M., Emilsson, L., Garborg, K., Rupinski, M., Dekker, E., Spaander, M., Bugajski, M., et al. (2022). Effect of Colonoscopy Screening on Risks of Colorectal Cancer and Related Death. N Engl J Med 387, 1547–1556. 10.1056/NEJMoa2208375.

7. Mazer, B.L., Lee, J.W., Roberts, N.J., Chu, L.C., Lennon, A.M., Klein, A.P., Eshleman, J.R., Fishman, E.K., Canto, M.I., Goggins, M.G., and Hruban, R.H. (2023). Screening for pancreatic cancer has the potential to save lives, but is it practical? Expert Rev Gastroenterol Hepatol 17, 555–574. 10.1080/17474124.2023.2217354.

8. Hur, C., Tramontano, A.C., Dowling, E.C., Brooks, G.A., Jeon, A., Brugge, W.R., Gazelle, G.S., Kong, C.Y., and Pandharipande, P.V. (2016). Early Pancreatic Ductal Adenocarcinoma Survival Is Dependent on Size: Positive Implications for Future Targeted Screening. Pancreas 45, 1062–1066. 10.1097/MPA.0000000000000587.

9. Egawa, S., Takeda, K., Fukuyama, S., Motoi, F., Sunamura, M., and Matsuno, S. (2004). Clinicopathological aspects of small pancreatic cancer. Pancreas 28, 235–240. 10.1097/00006676-200404000-00004.

10. Pollini, T., Wong, P., and Maker, A.V. (2023). The Landmark Series: Intraductal Papillary Mucinous Neoplasms of the Pancreas-From Prevalence to Early Cancer Detection. Ann Surg Oncol 30, 1453–1462. 10.1245/s10434-022-12870-w.

11. Choi, S.H., Park, S.H., Kim, K.W., Lee, J.Y., and Lee, S.S. (2017). Progression of Unresected Intraductal Papillary Mucinous Neoplasms of the Pancreas to Cancer: A Systematic Review and Meta-analysis. Clin Gastroenterol Hepatol 15, 1509–1520 e1504. 10.1016/j.cgh.2017.03.020.

12. Oyama, H., Tada, M., Takagi, K., Tateishi, K., Hamada, T., Nakai, Y., Hakuta, R., Ijichi, H., Ishigaki, K., Kanai, S., et al. (2020). Long-term Risk of Malignancy in Branch-Duct Intraductal Papillary Mucinous Neoplasms. Gastroenterology 158, 226–237 e225. 10.1053/j.gastro.2019.08.032.

13. Fischer, C.G., Beleva Guthrie, V., Braxton, A.M., Zheng, L., Wang, P., Song, Q., Griffin, J.F., Chianchiano, P.E., Hosoda, W., Niknafs, N., et al. (2019). Intraductal Papillary Mucinous Neoplasms Arise From Multiple Independent Clones, Each With Distinct Mutations. Gastroenterology 157, 1123–1137 e1122. 10.1053/j.gastro.2019.06.001.

14. Paolino, G., Esposito, I., Hong, S.M., Basturk, O., Mattiolo, P., Kaneko, T., Veronese, N., Scarpa, A., Adsay, V., and Luchini, C. (2022). Intraductal tubulopapillary neoplasm (ITPN) of the pancreas: a distinct entity among pancreatic tumors. Histopathology 81, 297–309. 10.1111/his.14698.

15. Laffan, T.A., Horton, K.M., Klein, A.P., Berlanstein, B., Siegelman, S.S., Kawamoto, S., Johnson, P.T., Fishman, E.K., and Hruban, R.H. (2008). Prevalence of unsuspected pancreatic cysts on MDCT. AJR Am J Roentgenol 191, 802–807. 10.2214/AJR.07.3340.

16. Wood, L.D., Adsay, N.V., Basturk, O., Brosens, L.A.A., Fukushima, N., Hong, S.M., Kim, S.J., Lee, J.W., Luchini, C., Noe, M., et al. (2023). Systematic review of challenging issues in pathology of intraductal papillary mucinous neoplasms. Pancreatology 23, 878–891. 10.1016/j.pan.2023.08.002.

17. Salvia, R., Burelli, A., Nepi, A., Caravati, A., Tomelleri, C., Dall’Olio, T., Casciani, F., Crino, S.F., Perri, G., and Marchegiani, G. (2023). Pancreatic cystic neoplasms: Still high rates of preoperative misdiagnosis in the guidelines and endoscopic ultrasound era. Surgery 174, 1410–1415. 10.1016/j.surg.2023.07.016.

18. Tanaka, M., Fernandez-del Castillo, C., Adsay, V., Chari, S., Falconi, M., Jang, J.Y., Kimura, W., Levy, P., Pitman, M.B., Schmidt, C.M., et al. (2012). International consensus guidelines 2012 for the management of IPMN and MCN of the pancreas. Pancreatology 12, 183–197. 10.1016/j.pan.2012.04.004.

19. Hasan, A., Visrodia, K., Farrell, J.J., and Gonda, T.A. (2019). Overview and comparison of guidelines for management of pancreatic cystic neoplasms. World J Gastroenterol 25, 4405–4413. 10.3748/wjg.v25.i31.4405.

20. Wilson, G.C., Maithel, S.K., Bentrem, D., Abbott, D.E., Weber, S., Cho, C., Martin, R.C., Scoggins, C.R., Kim, H.J., Merchant, N.B., et al. (2017). Are the Current Guidelines for the Surgical Management of Intraductal Papillary Mucinous Neoplasms of the Pancreas Adequate? A Multi-Institutional Study. J Am Coll Surg 224, 461–469. 10.1016/j.jamcollsurg.2016.12.031.

21. Srivastava, S., Koay, E.J., Borowsky, A.D., De Marzo, A.M., Ghosh, S., Wagner, P.D., and Kramer, B.S. (2019). Cancer overdiagnosis: a biological challenge and clinical dilemma. Nature Reviews Cancer 19, 349–358. 10.1038/s41568-019-0142-8.

22. Ciprani, D., Morales-Oyarvide, V., Qadan, M., Hank, T., Weniger, M., Harrison, J.M., Rodrigues, C., Horick, N.K., Mino-Kenudson, M., Ferrone, C.R., et al. (2020). An elevated CA 19-9 is associated with invasive cancer and worse survival in IPMN. Pancreatology 20, 729–735. 10.1016/j.pan.2020.04.002.

23. McIntyre, C.A., Rodrigues, C., Santharaman, A.V., Goldman, D.A., Javed, A.A., Ciprani, D., Pang, N., Lokshin, A., Gonen, M., Al Efishat, M.A., et al. (2022). Multiinstitutional Validation Study of Cyst Fluid Protein Biomarkers in Patients With Cystic Lesions of the Pancreas. Ann Surg 276, e129–e132. 10.1097/SLA.0000000000005314.

24. Levink, I.J.M., Jaarsma, S.C., Koopmann, B.D.M., van Riet, P.A., Overbeek, K.A., Meziani, J., Sprij, M., Casadei, R., Ingaldi, C., Polkowski, M., et al. (2023). The additive value of CA19.9 monitoring in a pancreatic cyst surveillance program. United European Gastroenterol J 11, 601–611. 10.1002/ueg2.12422.

25. Barutcuoglu, B., Oruc, N., Ak, G., Kucukokudan, S., Aydin, A., Nart, D., and Harman, M. (2022). Co-analysis of pancreatic cyst fluid carcinoembryonic antigen and glucose with novel cut-off levels better distinguishes between mucinous and non-mucinous neoplastic pancreatic cystic lesions. Ann Clin Biochem 59, 125–133. 10.1177/00045632211053998.

26. Carr, R.A., Yip-Schneider, M.T., Dolejs, S., Hancock, B.A., Wu, H., Radovich, M., and Schmidt, C.M. (2017). Pancreatic Cyst Fluid Vascular Endothelial Growth Factor A and Carcinoembryonic Antigen: A Highly Accurate Test for the Diagnosis of Serous Cystic Neoplasm. J Am Coll Surg. 10.1016/j.jamcollsurg.2017.05.003.

27. Pfluger, M.J., Jamouss, K.T., Afghani, E., Lim, S.J., Rodriguez Franco, S., Mayo, H., Spann, M., Wang, H., Singhi, A., Lennon, A.M., and Wood, L.D. (2023). Predictive ability of pancreatic cyst fluid biomarkers: A systematic review and meta-analysis. Pancreatology 23, 868–877. 10.1016/j.pan.2023.05.005.

28. Springer, S., Masica, D.L., Dal Molin, M., Douville, C., Thoburn, C.J., Afsari, B., Li, L., Cohen, J.D., Thompson, E., Allen, P.J., et al. (2019). A multimodality test to guide the management of patients with a pancreatic cyst. Sci Transl Med 11. 10.1126/scitranslmed.aav4772.

29. Kuboki, Y., Shimizu, K., Hatori, T., Yamamoto, M., Shibata, N., Shiratori, K., and Furukawa, T. (2015). Molecular biomarkers for progression of intraductal papillary mucinous neoplasm of the pancreas. Pancreas 44, 227–235. 10.1097/mpa.0000000000000253.

30. Li, Y., Lih, T.M., Dhanasekaran, S.M., Mannan, R., Chen, L., Cieslik, M., Wu, Y., Lu, R.J., Clark, D.J., Kolodziejczak, I., et al. (2023). Histopathologic and proteogenomic heterogeneity reveals features of clear cell renal cell carcinoma aggressiveness. Cancer Cell 41, 139–163 e117. 10.1016/j.ccell.2022.12.001.

31. Zhang, H., Liu, T., Zhang, Z., Payne, S.H., Zhang, B., McDermott, J.E., Zhou, J.Y., Petyuk, V.A., Chen, L., Ray, D., et al. (2016). Integrated Proteogenomic Characterization of Human High-Grade Serous Ovarian Cancer. Cell 166, 755–765. 10.1016/j.cell.2016.05.069.

32. Satpathy, S., Krug, K., Jean Beltran, P.M., Savage, S.R., Petralia, F., Kumar-Sinha, C., Dou, Y., Reva, B., Kane, M.H., Avanessian, S.C., et al. (2021). A proteogenomic portrait of lung squamous cell carcinoma. Cell 184, 4348–4371 e4340. 10.1016/j.cell.2021.07.016.

33. Cao, L., Huang, C., Cui Zhou, D., Hu, Y., Lih, T.M., Savage, S.R., Krug, K., Clark, D.J., Schnaubelt, M., Chen, L., et al. (2021). Proteogenomic characterization of pancreatic ductal adenocarcinoma. Cell 184, 5031–5052 e5026. 10.1016/j.cell.2021.08.023.

34. Clark, D.J., Dhanasekaran, S.M., Petralia, F., Pan, J., Song, X., Hu, Y., da Veiga Leprevost, F., Reva, B., Lih, T.M., Chang, H.Y., et al. (2019). Integrated Proteogenomic Characterization of Clear Cell Renal Cell Carcinoma. Cell 179, 964-983.e931. 10.1016/j.cell.2019.10.007.

35. Huang, C., Chen, L., Savage, S.R., Eguez, R.V., Dou, Y., Li, Y., da Veiga Leprevost, F., Jaehnig, E.J., Lei, J.T., Wen, B., et al. (2021). Proteogenomic insights into the biology and treatment of HPV-negative head and neck squamous cell carcinoma. Cancer Cell 39, 361-379.e316. 10.1016/j.ccell.2020.12.007.

36. Lih, T.M., Cho, K.C., Schnaubelt, M., Hu, Y., and Zhang, H. (2023). Integrated glycoproteomic characterization of clear cell renal cell carcinoma. Cell Rep 42, 112409. 10.1016/j.celrep.2023.112409.

37. Pan, J., Hu, Y., Sun, S., Chen, L., Schnaubelt, M., Clark, D., Ao, M., Zhang, Z., Chan, D., Qian, J., and Zhang, H. (2020). Glycoproteomics-based signatures for tumor subtyping and clinical outcome prediction of high-grade serous ovarian cancer. Nature Communications 11, 6139. 10.1038/s41467-020-19976-3.

38. Pinho, S.S., and Reis, C.A. (2015). Glycosylation in cancer: mechanisms and clinical implications. Nat Rev Cancer 15, 540–555. 10.1038/nrc3982.

39. Nikiforova, M.N., Wald, A.I., Spagnolo, D.M., Melan, M.A., Grupillo, M., Lai, Y.T., Brand, R.E., O’Broin-Lennon, A.M., McGrath, K., Park, W.G., et al. (2023). A Combined DNA/RNA-based Next-Generation Sequencing Platform to Improve the Classification of Pancreatic Cysts and Early Detection of Pancreatic Cancer Arising From Pancreatic Cysts. Ann Surg 278, e789–e797. 10.1097/SLA.0000000000005904.

40. Sohn, T.A., Yeo, C.J., Cameron, J.L., Hruban, R.H., Fukushima, N., Campbell, K.A., and Lillemoe, K.D. (2004). Intraductal papillary mucinous neoplasms of the pancreas: an updated experience. Ann Surg 239, 788-797; discussion 797-789. 10.1097/01.sla.0000128306.90650.aa.

41. Valsangkar, N.P., Morales-Oyarvide, V., Thayer, S.P., Ferrone, C.R., Wargo, J.A., Warshaw, A.L., and Fernandez-del Castillo, C. (2012). 851 resected cystic tumors of the pancreas: a 33-year experience at the Massachusetts General Hospital. Surgery 152, S4–12. 10.1016/j.surg.2012.05.033.

42. Springer, S., Wang, Y., Dal Molin, M., Masica, D.L., Jiao, Y., Kinde, I., Blackford, A., Raman, S.P., Wolfgang, C.L., Tomita, T., et al. (2015). A combination of molecular markers and clinical features improve the classification of pancreatic cysts. Gastroenterology 149, 1501–1510. 10.1053/j.gastro.2015.07.041.

43. Wu, J., Jiao, Y., Dal Molin, M., Maitra, A., de Wilde, R.F., Wood, L.D., Eshleman, J.R., Goggins, M.G., Wolfgang, C.L., Canto, M.I., et al. (2011). Whole-exome sequencing of neoplastic cysts of the pancreas reveals recurrent mutations in components of ubiquitin-dependent pathways. Proc Natl Acad Sci U S A 108, 21188–21193. 10.1073/pnas.1118046108.

44. Gillet, L.C., Navarro, P., Tate, S., Röst, H., Selevsek, N., Reiter, L., Bonner, R., and Aebersold, R. (2012). Targeted data extraction of the MS/MS spectra generated by data-independent acquisition: a new concept for consistent and accurate proteome analysis. Mol Cell Proteomics 11, O111.016717. 10.1074/mcp.O111.016717.

45. Meier, F., Brunner, A.D., Frank, M., Ha, A., Bludau, I., Voytik, E., Kaspar-Schoenefeld, S., Lubeck, M., Raether, O., Bache, N., et al. (2020). diaPASEF: parallel accumulation-serial fragmentation combined with data-independent acquisition. Nat Methods 17, 1229–1236. 10.1038/s41592-020-00998-0.

46. Bruderer, R., Bernhardt, O.M., Gandhi, T., Miladinović, S.M., Cheng, L.Y., Messner, S., Ehrenberger, T., Zanotelli, V., Butscheid, Y., Escher, C., et al. (2015). Extending the limits of quantitative proteome profiling with data-independent acquisition and application to acetaminophen-treated three-dimensional liver microtissues. Mol Cell Proteomics 14, 1400–1410. 10.1074/mcp.M114.044305.

47. Furukawa, T. (2022). Intraductal Neoplasms of the Pancreas. In The IASGO Textbook of Multi-Disciplinary Management of Hepato-Pancreato-Biliary Diseases, M. Makuuchi, N. Kokudo, I. Popescu, J. Belghiti, H.-S. Han, K. Takaori, and D.G. Duda, eds. (Springer Nature Singapore), pp. 77–84. 10.1007/978-981-19-0063-1_10.

48. Nagata, K., Horinouchi, M., Saitou, M., Higashi, M., Nomoto, M., Goto, M., and Yonezawa, S. (2007). Mucin expression profile in pancreatic cancer and the precursor lesions. J Hepatobiliary Pancreat Surg 14, 243–254. 10.1007/s00534-006-1169-2.

49. Kaur, S., Kumar, S., Momi, N., Sasson, A.R., and Batra, S.K. (2013). Mucins in pancreatic cancer and its microenvironment. Nature Reviews Gastroenterology & Hepatology 10, 607–620. 10.1038/nrgastro.2013.120.

50. Gautam, S.K., Khan, P., Natarajan, G., Atri, P., Aithal, A., Ganti, A.K., Batra, S.K., Nasser, M.W., and Jain, M. (2023). Mucins as Potential Biomarkers for Early Detection of Cancer. Cancers 15, 1640.

51. Pollini, T., Adsay, V., Capurso, G., Dal Molin, M., Esposito, I., Hruban, R., Luchini, C., Maggino, L., Matthaei, H., Marchegiani, G., et al. (2022). The tumour immune microenvironment and microbiome of pancreatic intraductal papillary mucinous neoplasms. Lancet Gastroenterol Hepatol 7, 1141–1150. 10.1016/S2468-1253(22)00235-7.

52. Iacobuzio-Donahue, C.A., Maitra, A., Olsen, M., Lowe, A.W., van Heek, N.T., Rosty, C., Walter, K., Sato, N., Parker, A., Ashfaq, R., et al. (2003). Exploration of global gene expression patterns in pancreatic adenocarcinoma using cDNA microarrays. Am J Pathol 162, 1151–1162. 10.1016/s0002-9440(10)63911-9.

53. Iacobuzio-Donahue, C.A., Maitra, A., Shen-Ong, G.L., van Heek, T., Ashfaq, R., Meyer, R., Walter, K., Berg, K., Hollingsworth, M.A., Cameron, J.L., et al. (2002). Discovery of novel tumor markers of pancreatic cancer using global gene expression technology. Am J Pathol 160, 1239–1249. 10.1016/s0002-9440(10)62551-5.

54. Li, Y., Chang, R.B., Stone, M.L., Delman, D., Markowitz, K., Xue, Y., Coho, H., Herrera, V.M., Li, J.H., Zhang, L., et al. (2024). Multimodal immune phenotyping reveals microbial-T cell interactions that shape pancreatic cancer. Cell Rep Med 5, 101397. 10.1016/j.xcrm.2024.101397.

55. Turula, H., and Wobus, C.E. (2018). The Role of the Polymeric Immunoglobulin Receptor and Secretory Immunoglobulins during Mucosal Infection and Immunity. Viruses 10. 10.3390/v10050237.

56. Ohtsuka, T., Fernandez-Del Castillo, C., Furukawa, T., Hijioka, S., Jang, J.Y., Lennon, A.M., Miyasaka, Y., Ohno, E., Salvia, R., Wolfgang, C.L., and Wood, L.D. (2024). International evidence-based Kyoto guidelines for the management of intraductal papillary mucinous neoplasm of the pancreas. Pancreatology 24, 255–270. 10.1016/j.pan.2023.12.009.

57. European evidence-based guidelines on pancreatic cystic neoplasms. (2018). Gut 67, 789–804. 10.1136/gutjnl-2018-316027.

58. Büll, C., Stoel, M.A., den Brok, M.H., and Adema, G.J. (2014). Sialic acids sweeten a tumor’s life. Cancer Res 74, 3199–3204. 10.1158/0008-5472.Can-14-0728.

59. Ahmed, M., Biswas, T., and Mondal, S. (2023). The strategic involvement of IRS in cancer progression. Biochem Biophys Res Commun 680, 141–160. 10.1016/j.bbrc.2023.09.036.

60. Ferreira, I.G., Carrascal, M., Mineiro, A.G., Bugalho, A., Borralho, P., Silva, Z., Dall’olio, F., and Videira, P.A. (2019). Carcinoembryonic antigen is a sialyl Lewis x/a carrier and an EIZselectin ligand in nonIZsmall cell lung cancer. Int J Oncol 55, 1033–1048. 10.3892/ijo.2019.4886.

61. Gao, H.F., Wang, Q.Y., Zhang, K., Chen, L.Y., Cheng, C.S., Chen, H., Meng, Z.Q., Zhou, S.M., and Chen, Z. (2019). Overexpressed N-fucosylation on the cell surface driven by FUT3, 5, and 6 promotes cell motilities in metastatic pancreatic cancer cell lines. Biochem Biophys Res Commun 511, 482–489. 10.1016/j.bbrc.2019.02.092.

62. Zhou, H., Zhao, J., Yang, X., Liu, J., and Huang, W. (2022). Study on the Expression of beta-1,3-N-acetylglucosaminyltransferase 3 in Gastric Cancer and the Mechanism Promoting Gastric Cancer Progression Based on the Extraction Method of Nanomagnetic Beads. J Biomed Nanotechnol 18, 677–692. 10.1166/jbn.2022.3296.

63. Zhang, J., Tian, Y., Mo, S., and Fu, X. (2022). Overexpressing PLOD Family Genes Predict Poor Prognosis in Pancreatic Cancer. Int J Gen Med 15, 3077–3096. 10.2147/ijgm.S341332.

64. Seifert, A.M., Reiche, C., Heiduk, M., Tannert, A., Meinecke, A.C., Baier, S., von Renesse, J., Kahlert, C., Distler, M., Welsch, T., et al. (2020). Detection of pancreatic ductal adenocarcinoma with galectin-9 serum levels. Oncogene 39, 3102–3113. 10.1038/s41388-020-1186-7.

65. Omori, Y., Ono, Y., Kobayashi, T., Motoi, F., Karasaki, H., Mizukami, Y., Makino, N., Ueno, Y., Unno, M., and Furukawa, T. (2020). How does intestinal-type intraductal papillary mucinous neoplasm emerge? CDX2 plays a critical role in the process of intestinal differentiation and progression. Virchows Arch 477, 21–31. 10.1007/s00428-020-02806-8.

66. Pan, S., Brand, R.E., Lai, L.A., Dawson, D.W., Donahue, T.R., Kim, S., Khalaf, N.I., Othman, M.O., Fisher, W.E., Bronner, M.P., et al. (2021). Proteome heterogeneity and malignancy detection in pancreatic cyst fluids. Clin Transl Med 11, e506. 10.1002/ctm2.506.

67. Sawai, H., Okada, Y., Tanaka, M., Funahashi, H., Hayakawa, T., and Manabe, T. (2004). Expression of integrins in intraductal papillary-mucinous tumors of the pancreas as an indicator of malignancy. Pancreas 28, 20–24. 10.1097/00006676-200401000-00003.

68. Zhu, H., Wang, G., Zhu, H., and Xu, A. (2021). ITGA5 is a prognostic biomarker and correlated with immune infiltration in gastrointestinal tumors. BMC Cancer 21, 269. 10.1186/s12885-021-07996-1.

69. Ishida, M., Egawa, S., Aoki, T., Sakata, N., Mikami, Y., Motoi, F., Abe, T., Fukuyama, S., Sunamura, M., Unno, M., et al. (2007). Characteristic clinicopathological features of the types of intraductal papillary-mucinous neoplasms of the pancreas. Pancreas 35, 348–352. 10.1097/mpa.0b013e31806da090.

70. Getu, A.A., Tigabu, A., Zhou, M., Lu, J., Fodstad, Ø., and Tan, M. (2023). New frontiers in immune checkpoint B7-H3 (CD276) research and drug development. Molecular Cancer 22, 43. 10.1186/s12943-023-01751-9.

71. Zhou, W.T., and Jin, W.L. (2021). B7-H3/CD276: An Emerging Cancer Immunotherapy. Front Immunol 12, 701006. 10.3389/fimmu.2021.701006.

72. Michaels, A.D., Newhook, T.E., Adair, S.J., Morioka, S., Goudreau, B.J., Nagdas, S., Mullen, M.G., Persily, J.B., Bullock, T.N.J., Slingluff, C.L., Jr., et al. (2018). CD47 Blockade as an Adjuvant Immunotherapy for Resectable Pancreatic Cancer. Clin Cancer Res 24, 1415–1425. 10.1158/1078-0432.Ccr-17-2283.

73. Hayat, S.M.G., Bianconi, V., Pirro, M., Jaafari, M.R., Hatamipour, M., and Sahebkar, A. (2020). CD47: role in the immune system and application to cancer therapy. Cellular Oncology 43, 19–30. 10.1007/s13402-019-00469-5.

74. Logtenberg, M.E.W., Scheeren, F.A., and Schumacher, T.N. (2020). The CD47-SIRPα Immune Checkpoint. Immunity 52, 742–752. 10.1016/j.immuni.2020.04.011.

75. Theocharis, A.D., Skandalis, S.S., Gialeli, C., and Karamanos, N.K. (2016). Extracellular matrix structure. Adv Drug Deliv Rev 97, 4–27. 10.1016/j.addr.2015.11.001.

76. Barczyk, M., Carracedo, S., and Gullberg, D. (2010). Integrins. Cell and Tissue Research 339, 269–280. 10.1007/s00441-009-0834-6.

77. Domogatskaya, A., Rodin, S., and Tryggvason, K. (2012). Functional diversity of laminins. Annu Rev Cell Dev Biol 28, 523–553. 10.1146/annurev-cellbio-101011-155750.

78. Moremen, K.W., Tiemeyer, M., and Nairn, A.V. (2012). Vertebrate protein glycosylation: diversity, synthesis and function. Nat Rev Mol Cell Biol 13, 448–462. 10.1038/nrm3383.

79. Ribatti, D., Tamma, R., and Annese, T. (2020). Epithelial-Mesenchymal Transition in Cancer: A Historical Overview. Transl Oncol 13, 100773. 10.1016/j.tranon.2020.100773.

80. Light, A., and Janska, H. (1989). Enterokinase (enteropeptidase): comparative aspects. Trends Biochem Sci 14, 110–112. 10.1016/0968-0004(89)90133-3.

81. Jönsson, P., Källén, R., Montgomery, A., and Borgström, A. (1995). Protease activation in the porcine pancreatic allograft during preservation. Pancreas 11, 256–260. 10.1097/00006676-199510000-00007.

82. Kylänpää-Bäck, M.L., Kemppainen, E., and Puolakkainen, P. (2002). Trypsin-based laboratory methods and carboxypeptidase activation peptide in acute pancreatitis. Jop 3, 34–48.

83. Assarzadegan, N., Thompson, E., Salimian, K., Gaida, M.M., Brosens, L.A.A., Wood, L., Ali, S.Z., and Hruban, R.H. (2021). Pathology of intraductal papillary mucinous neoplasms. Langenbecks Arch Surg 406, 2643–2655. 10.1007/s00423-021-02201-0.

84. Noë, M., Niknafs, N., Fischer, C.G., Hackeng, W.M., Beleva Guthrie, V., Hosoda, W., Debeljak, M., Papp, E., Adleff, V., White, J.R., et al. (2020). Genomic characterization of malignant progression in neoplastic pancreatic cysts. Nat Commun 11, 4085. 10.1038/s41467-020-17917-8.

85. Li, M.X., Wang, H.Y., Yuan, C.H., Ma, Z.L., Jiang, B., Li, L., Zhang, L., and Xiu, D.R. (2021). Establishment of a Macrophage Phenotypic Switch Related Prognostic Signature in Patients With Pancreatic Cancer. Front Oncol 11, 619517. 10.3389/fonc.2021.619517.

86. Furukawa, T., Klöppel, G., Volkan Adsay, N., Albores-Saavedra, J., Fukushima, N., Horii, A., Hruban, R.H., Kato, Y., Klimstra, D.S., Longnecker, D.S., et al. (2005). Classification of types of intraductal papillary-mucinous neoplasm of the pancreas: a consensus study. Virchows Arch 447, 794–799. 10.1007/s00428-005-0039-7.

87. Adsay, V., Mino-Kenudson, M., Furukawa, T., Basturk, O., Zamboni, G., Marchegiani, G., Bassi, C., Salvia, R., Malleo, G., Paiella, S., et al. (2016). Pathologic Evaluation and Reporting of Intraductal Papillary Mucinous Neoplasms of the Pancreas and Other Tumoral Intraepithelial Neoplasms of Pancreatobiliary Tract: Recommendations of Verona Consensus Meeting. Ann Surg 263, 162–177. 10.1097/sla.0000000000001173.

88. Gaujoux, R., and Seoighe, C. (2010). A flexible R package for nonnegative matrix factorization. BMC Bioinformatics 11, 367. 10.1186/1471-2105-11-367.

89. Aran, D., Hu, Z., and Butte, A.J. (2017). xCell: digitally portraying the tissue cellular heterogeneity landscape. Genome Biol 18, 220. 10.1186/s13059-017-1349-1.

90. Wherry, E.J. (2011). T cell exhaustion. Nat Immunol 12, 492–499. 10.1038/ni.2035.

91. Goronzy, J.J., and Weyand, C.M. (2019). Mechanisms underlying T cell ageing. Nat Rev Immunol 19, 573–583. 10.1038/s41577-019-0180-1.

92. Mukherji, R., Debnath, D., Hartley, M.L., and Noel, M.S. (2022). The Role of Immunotherapy in Pancreatic Cancer. Curr Oncol 29, 6864–6892. 10.3390/curroncol29100541.

93. Pishvaian, M.J., Blais, E.M., Brody, J.R., Lyons, E., DeArbeloa, P., Hendifar, A., Mikhail, S., Chung, V., Sahai, V., Sohal, D.P.S., et al. (2020). Overall survival in patients with pancreatic cancer receiving matched therapies following molecular profiling: a retrospective analysis of the Know Your Tumor registry trial. Lancet Oncol 21, 508–518. 10.1016/s1470-2045(20)30074-7.

94. Khoury, R.E., Kabir, C., Maker, V.K., Banulescu, M., Wasserman, M., and Maker, A.V. (2018). What is the Incidence of Malignancy in Resected Intraductal Papillary Mucinous Neoplasms? An Analysis of Over 100 US Institutions in a Single Year. Ann Surg Oncol 25, 1746–1751. 10.1245/s10434-018-6425-6.

95. Kawada, N., Uehara, H., Nagata, S., Tsuchishima, M., Tsutsumi, M., and Tomita, Y. (2016). Mural nodule of 10 mm or larger as predictor of malignancy for intraductal papillary mucinous neoplasm of the pancreas: Pathological and radiological evaluations. Pancreatology 16, 441–448. 10.1016/j.pan.2015.12.008.

96. Zhao, W., Liu, S., Cong, L., and Zhao, Y. (2022). Imaging Features for Predicting High-Grade Dysplasia or Malignancy in Branch Duct Type Intraductal Papillary Mucinous Neoplasm of the Pancreas: A Systematic Review and Meta-Analysis. Ann Surg Oncol 29, 1297–1312. 10.1245/s10434-021-10662-2.

97. Basturk, O., Hong, S.M., Wood, L.D., Adsay, N.V., Albores-Saavedra, J., Biankin, A.V., Brosens, L.A., Fukushima, N., Goggins, M., Hruban, R.H., et al. (2015). A Revised Classification System and Recommendations From the Baltimore Consensus Meeting for Neoplastic Precursor Lesions in the Pancreas. Am J Surg Pathol 39, 1730–1741. 10.1097/pas.0000000000000533.

98. Rezaee, N., Barbon, C., Zaki, A., He, J., Salman, B., Hruban, R.H., Cameron, J.L., Herman, J.M., Ahuja, N., Lennon, A.M., et al. (2016). Intraductal papillary mucinous neoplasm (IPMN) with high-grade dysplasia is a risk factor for the subsequent development of pancreatic ductal adenocarcinoma. HPB (Oxford) 18, 236–246. 10.1016/j.hpb.2015.10.010.

99. Maitra, A., Iacobuzio-Donahue, C., Rahman, A., Sohn, T.A., Argani, P., Meyer, R., Yeo, C.J., Cameron, J.L., Goggins, M., Kern, S.E., et al. (2002). Immunohistochemical validation of a novel epithelial and a novel stromal marker of pancreatic ductal adenocarcinoma identified by global expression microarrays: sea urchin fascin homolog and heat shock protein 47. Am J Clin Pathol 118, 52–59. 10.1309/3pam-p5wl-2lv0-r4eg.

100. Iacobuzio-Donahue, C.A., Ryu, B., Hruban, R.H., and Kern, S.E. (2002). Exploring the host desmoplastic response to pancreatic carcinoma: gene expression of stromal and neoplastic cells at the site of primary invasion. Am J Pathol 160, 91–99. 10.1016/s0002-9440(10)64353-2.

101. Castellano, E., and Downward, J. (2011). RAS Interaction with PI3K: More Than Just Another Effector Pathway. Genes Cancer 2, 261–274. 10.1177/1947601911408079.

102. Mertins, P., Tang, L.C., Krug, K., Clark, D.J., Gritsenko, M.A., Chen, L., Clauser, K.R., Clauss, T.R., Shah, P., Gillette, M.A., et al. (2018). Reproducible workflow for multiplexed deep-scale proteome and phosphoproteome analysis of tumor tissues by liquid chromatography–mass spectrometry. Nature Protocols 13, 1632–1661. 10.1038/s41596-018-0006-9.

103. Liao, Y., Wang, J., Jaehnig, E.J., Shi, Z., and Zhang, B. (2019). WebGestalt 2019: gene set analysis toolkit with revamped UIs and APIs. Nucleic Acids Res 47, W199–w205. 10.1093/nar/gkz401.

104. Braxton, A.M., Kiemen, A.L., Grahn, M.P., Forjaz, A., Parksong, J., Mahesh Babu, J., Lai, J., Zheng, L., Niknafs, N., Jiang, L., et al. (2024). 3D genomic mapping reveals multifocality of human pancreatic precancers. Nature 629, 679–687. 10.1038/s41586-024-07359-3.

